# Sobetirome, a thyroid hormone receptor beta agonist, is a potential therapeutic agent for pulmonary fibrosis

**DOI:** 10.64898/2026.08.31.747360

**Authors:** Thomas Bärnthaler, Shuizi Ding, Taylor S. Adams, Kady-Ann Rose, Reina Rangel, Johad Khoury, Melika Salimi, Sabina Anderson, Liqin Lin, Karen Velasco Alzate, Fernando Poli De Frias, Giuseppe Deluliis, Fadi Nicola, Carlos Cosme, Aurelien Justet, Xinran Liu, Ivan O. Rosas, Naftali Kaminski, Farida Ahangari

## Abstract

Idiopathic pulmonary fibrosis (IPF) is a progressive and fatal disease with limited treatment options. Our group previously identified the antifibrotic potential of thyroid hormone, triiodothyronine (T3); however, clinical translation of thyroid hormone therapy is limited by its systemic adverse effects. In this study, we investigate whether sobetirome, a selective and well-tolerated thyroid hormone receptor beta (THRB) agonist, offers antifibrotic benefits of thyroid hormone while minimizing systemic toxicity.

Our study reveals that sobetirome, administered via intraperitoneal or inhalational routes, effectively mitigates bleomycin-induced pulmonary fibrosis in mice, with no evidence of toxicity. We identified that sobetirome restores mitochondrial homeostasis via activating the THRB–PPARGC1α axis. This protects alveolar type II epithelial cells from injury-induced apoptosis while selectively inducing apoptosis and metabolic reprogramming in apoptosis-resistant IPF fibroblasts. Cell-specific deletion of *Ppargc1α* in either alveolar epithelial cells or fibroblasts abolishes sobetirome-mediated protection, establishing PPARGC1α as an essential mediator of therapeutic response. Importantly, sobetirome reverses fibrosis-associated transcriptional programs in human IPF lung tissue, reducing expression of key fibrosis-associated genes, including collagen I alpha 1 (*COL1A1*), collagen III alpha 1 (*COL3A1*), periostin (*POSTN),* cathepsin K (*CTSK*), and Chitinase 3 Like 1 (*CHI3L1*), while promoting extracellular matrix remodeling, epithelial restoration, and tissue homeostasis.

Collectively, our findings identify THRB activation as a novel metabolic strategy for reversing pulmonary fibrosis. Across complementary *in vitro*, *in vivo*, and human *ex vivo* models, sobetirome restores mitochondrial function, modulates apoptotic pathways in pathogenic cells, and promotes fibrosis resolution, highlighting its potential as a lung-targeted therapeutic approach for IPF and other fibrotic lung diseases.

**One sentence summary:** Sobetirome, a thyroid hormone receptor beta agonist, exerts potent antifibrotic effects in preclinical models of pulmonary fibrosis across *in vitro*, *in vivo*, and *ex vivo* settings by acting on both lung epithelial and fibroblast cells.

## Introduction

Idiopathic pulmonary fibrosis (IPF) is a chronic progressive disease characterized by respiratory insufficiency, often necessitating lung transplantation [1]. Without intervention, the disease is typically fatal, with a median survival of 3–5 years [2]. Although the three approved drugs, pirfenidone, nintedanib, and nerandomilast have shown the ability to slow disease progression and improve clinical outcomes, none are curative, and they do not reverse established fibrosis or restore the damaged alveolar architecture [2–6]. There remains a critical need for the continued development of more effective therapeutic strategies.

The primary hallmark of IPF is the progressive loss of alveoli, the functional unit of gas exchange, and their replacement with fibrotic tissue. This process is marked by increased apoptosis and a significant reduction in alveolar epithelial cells, particularly type II cells (ATIIs) [7] while fibroblasts are differentiating to myofibroblasts and are likely to become resistant to apoptosis [8, 9]. These seemingly contradictory effects have been termed the “apoptosis paradox” in IPF and are believed to contribute to disease progression and pathogenesis [8]. Importantly, mouse models of lung fibrosis have shown that both increased alveolar epithelial cell apoptosis and reduced apoptosis in fibroblasts can promote lung fibrosis [10, 11]. Consequently, it has been proposed that drugs that can impact both cell types would show greater promise as a potential therapeutic agent [12].

In recent years, increasing evidence has highlighted the critical role of mitochondrial dysfunction and metabolic reprogramming in the pathogenesis of IPF. Increased extracellular mitochondrial DNA has been recognized as an independent predictor of poor survival in IPF and is accompanied by marked metabolic reprogramming in IPF fibroblasts, characterized by a preferential reliance on aerobic glycolysis over oxidative phosphorylation [13]. Notably, IPF myofibroblasts display fragmented mitochondria, impaired mitochondrial function, and resistance to apoptosis. Pharmacological AMP-activated protein kinase (AMPK) activation using AICAR restores mitochondrial function and apoptosis sensitivity in these cells. Additionally, AMPK deficiency alters mitochondrial function, reduces basal oxygen consumption and peak respiratory capacity [14]. In parallel, ATII cells in IPF exhibit profound mitochondrial dysfunction characterized by defective mitophagy, accumulation of damaged mitochondria, and increased apoptosis [15]. Loss of PTEN-induced kinase 1 (*PINK1*), a key regulator of mitophagy, disrupts mitochondrial homeostasis and promotes pulmonary fibrosis, while accumulation of mitochondrial damage-associated signals, including extracellular mitochondrial DNA (mtDNA), enhances profibrotic responses in lung epithelial cells [16]. Consistent with these findings, TGF-β1 induces epithelial mitochondrial dysfunction and alters *PINK1*-dependent mitochondrial pathways, contributing to fibrotic progression [17]. Furthermore, we demonstrated that ATII cells from bleomycin-injured lungs exhibit impaired mitochondrial bioenergetics [18].

Thyroid hormones (most notably triiodothyronine (T3) and thyroxine (T4)) have long been recognized as crucial regulators of metabolism in mammals. *In vivo*, T4 is converted to T3, which is considered the biologically more active form via iodothyronine deiodinase (DIO2) and binds to either thyroid hormone receptor alpha (THRA) or beta (THRB) [19]. While similar in their effects, there are considerable differences in tissue expression between these receptors, with predominantly THRB expression in the lung and THRA in skeletal or cardiac muscle [19, 20]. Importantly, there is strong evidence that many of their effects are at least partly mediated via mitochondria [21]. For example, T3 has been shown to increase mitobiogenesis, basal respiration, and proton leak *in vitro* and *in vivo* [21–23]. Many of these effects depend upon peroxisome proliferator-activated receptor gamma coactivator 1α (PPARGC1α or PGC-1α), which is considered the master regulator of mitobiogenesis [24]. We previously demonstrated that thyroid hormone agonism exerts anti-fibrotic effects, mainly via the protection of epithelial cells from apoptosis and restoration of their mitochondrial function [18]. However, there are certain limitations in using T3 or T4 in patients; first and foremost, the increased metabolic rate goes hand in hand with potential thyrotoxic side effects, such as weight loss, tachycardia, or arrhythmia [25]. Sobetirome is a thyroid hormone agonist that shows specificity for the THRB over the THRA isoform and hence lacks many of the side effects of T3 or T4, foremost the cardiotoxicity, and has also been tested with a favorable safety profile in human studies [26]. Building on these unique properties, our group was the first to establish that sobetirome effectively blunts bleomycin-induced pulmonary fibrosis in mice [18]. Additionally, a more recent work identified thyroid hormone signaling as a key regulator of lung regeneration and demonstrated that THRB activation by sobetirome promotes alveolar epithelial repair through modulation of the ATII to ATI differentiation [27].

In the present study, we comprehensively evaluated the therapeutic and cellular effects of sobetirome across complementary models of pulmonary fibrosis, including murine models, primary human and murine cells, and human lung tissue. We specifically investigated its effects on fibroblast activation and alveolar epithelial repair to define its main cellular targets and elucidate the molecular mechanisms underlying its antifibrotic activity. These studies provide mechanistic and translational insight into the therapeutic potential of selective THRB activation for the treatment of IPF.

## Results

### Intraperitoneal Administration of Sobetirome Ameliorates Pulmonary Fibrosis in the Bleomycin-Induced Mouse Model

To confirm the therapeutic potential of sobetirome, we assessed its effects in a bleomycin-induced pulmonary fibrosis mouse model. We initially performed dose-ranging studies to determine the optimal therapeutic dose of sobetirome. To this end, we tested various doses of sobetirome (10, 30, 100, 300, and 1000 microgram/kilogram (ug/kg) body weight) administered intraperitoneally (IP) every other day for 14 days in wild-type (WT) mice. The goal was to determine the optimally effective dose while ensuring safety in mice. At the end of the study, we collected lung tissues, blood, and bronchoalveolar lavage (BAL) fluids to assess liver function and blood lipid levels, as well as to evaluate the expression levels of *Ppargc1a*, a well-established target gene of sobetirome and its potential receptor *Thrb*. Based on these preliminary experiments, we selected 300 µg/kg as the optimal dose for our *in vivo* study, as it effectively increased *Ppargc1a* and *Thrb* expression and decreased LDL levels without inducing liver toxicity (**Figure S1A-E)**.

Next, we assessed the antifibrotic effects of sobetirome at the selected dose in the standard bleomycin-induced lung fibrosis model **(Fig. 1A)**. As expected, bleomycin-treated mice exhibited significant weight loss by day 8 compared with saline-treated controls. In contrast, treatment with sobetirome significantly reversed this effect, leading to a marked increase in body weight by day 21 **(Fig. 1B)**. Importantly, sobetirome alone had no significant impact on body weight relative to vehicle-treated mice. Histological analysis and hydroxyproline content measurements of these lungs demonstrated that sobetirome significantly reduced lung fibrosis in the bleomycin model **(Fig. 1C, D**, and **E**). Lung function analysis using the FlexiVent system demonstrated that bleomycin administration induced marked pulmonary dysfunction, as evidenced by significant reductions in static compliance and inspiratory capacity **(Fig. 1F** and **G**). Consistent with improvements observed in other parameters, sobetirome treatment substantially attenuated these deficits, restoring lung mechanics toward control levels **(Fig. 1F** and **G**) and similarly improving pressure–volume relationships **(Fig. 1H)**. Moreover, gene expression analysis of lung tissue revealed that sobetirome suppressed bleomycin-induced profibrotic gene expression, including *Col1a1* (collagen type I a 1) **(Fig. 1I**).

**Figure 1.**
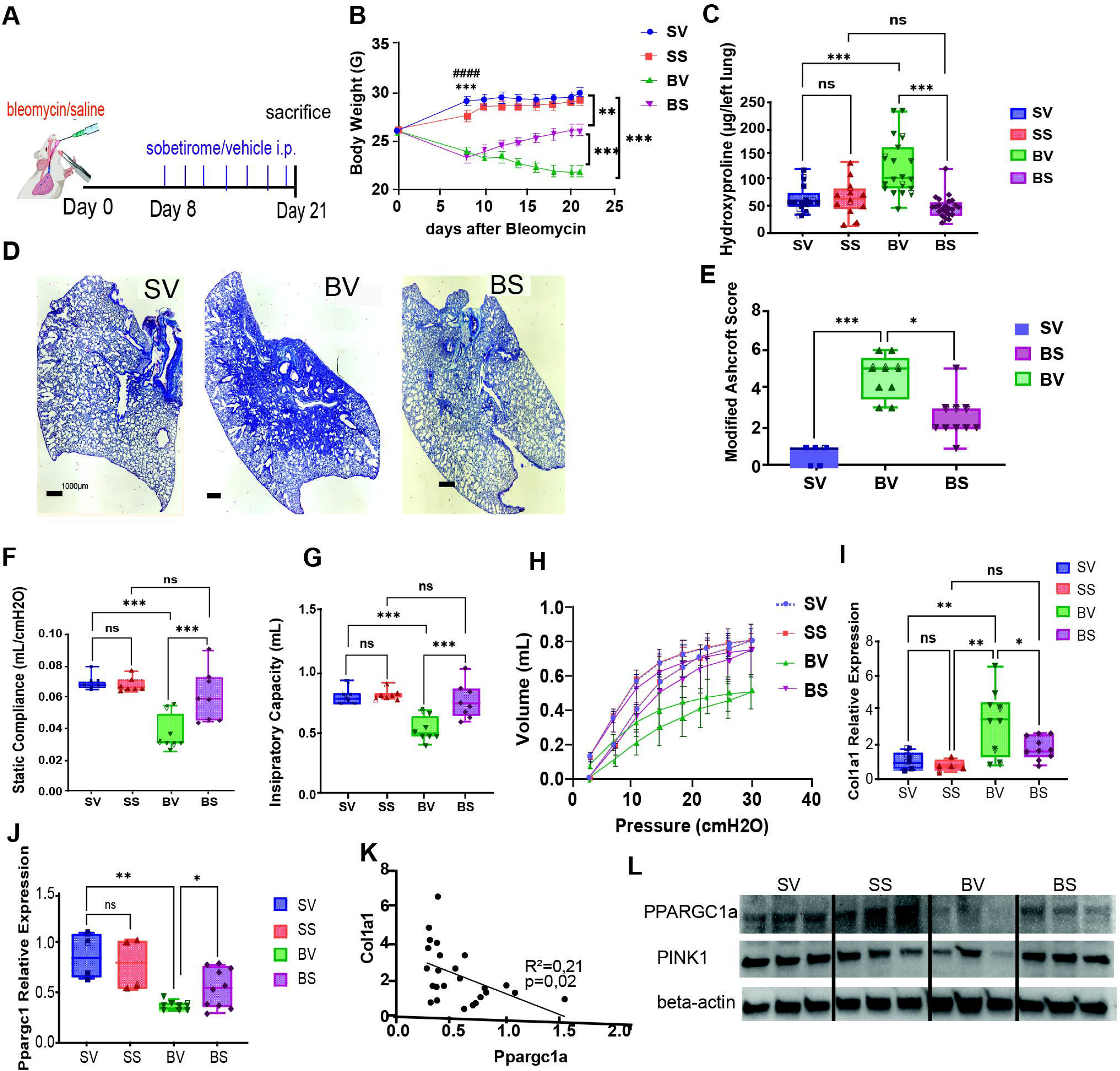
Intraperitoneal administration of sobetirome ameliorates pulmonary fibrosis in a mouse model. (**A**) Experimental Design. Mice were divided into four groups: 1-Saline + Vehicle (SV), 2-Saline + Sobetirome (SS),3-Bleomycin (1,5U/kg) + Vehicle (BV), 4-Bleomycin + Sobetirome (BS). IP administration of 300 mg/kg sobetirome was performed every other day. (**B**) Total Body Weight Changes. Weights were measured every other day, starting at the beginning of sobetirome/vehicle treatment, and the last observation was carried forward for deceased mice. (**C**) Hydroxyproline levels, as a measurement for lung collagen, were assayed from lung lysates. **(D)** Representative Masson trichrome-stained lung sections showing collagen deposition. Scale bars = 1 mm. Histological sections were evaluated in a blinded manner. **(E)** Quantification of pulmonary fibrosis using the modified Ashcroft score. **(F–H)** Lung function assessment using the FlexiVent system: **(F)** static compliance, **(G)** total lung capacity, and **(H)** pressure–volume curves. **(I–K)** Gene expression analysis of whole lung tissue by qPCR: **(I)** *Col1a1*, **(J)** *Ppargc1α*, and **(K)** linear regression analysis showing the correlation between *Col1a1* and *Ppargc1α* expression. **(L)** Representative immunoblot showing the indicated protein targets across experimental groups. Statistical analyses were performed using one-way ANOVA for panels C, F, G, I, and J; the Kruskal–Wallis test for panel E; and two-way ANOVA followed by Holm-Sidak multiple-comparisons testing for panels B and H. Data are presented from n=12–23 animals for histological and molecular analyses and n=7–8 animals for lung function measurements, respectively. *=p<0.05, **=p<0.01, ***=p<0.001

In line with the observed improvements in lung mechanics and attenuation of fibrotic gene expression, bleomycin treatment markedly suppressed *Ppargc1a* expression, an effect that was significantly restored by sobetirome (**Fig. 1J)**. Notably, *Ppargc1a* expression levels inversely correlated with *Col1a1* expression in the lungs of these animals (R² = 0.21, *p* = 0.02) **(Fig. 1K)**, suggesting a potential mechanistic link between metabolic regulation and fibrotic remodeling. Consistent with these transcriptional changes, sobetirome treatment increased both Ppargc1a and Pink1 protein levels after bleomycin administration, paralleling previously reported effects of T3 **(Fig. 1L)** [18].

Collectively, these findings demonstrate that sobetirome markedly attenuates bleomycin-induced pulmonary fibrosis, implicating a central role for metabolic regulation in its antifibrotic effects and underscoring this agent as a promising therapeutic candidate for fibrotic lung disease.

### Sobetirome Exerts Minimal Effects in Healthy Lungs but Reverses Fibrosis-Associated Gene Signatures in Mice after Bleomycin

To explore more on the antifibrotic effect of sobetirome in the bleomycin murine model, we performed bulk RNA sequencing on the lung tissues harvested from all four groups of mice: (Saline + Vehicle, Saline + Sobetirome, Bleomycin + Vehicle, and Bleomycin + Sobetirome). Principal component analysis (PCA) shows relatively minimal variance between Sobetirome-treated and nontreated samples in the absence of bleomycin (**Fig.2A**), indicating that the effects of sobetirome are dependent on the condition of injury and confirming the safety aspect of sobetirome in a healthy lung. Leveraging the experimental multifactor design, we tested for the main effects of the bleomycin condition, the sobetirome treatment, and the interaction effect between the two (**Fig.2B**). Consistent with the PCA results, the main effect of Sobetirome treatment alone is relatively muted: the 21 differentially effected genes comprise only 0.2% of the total differentially effected genes across tests (**Fig.2C**). In contrast, the interaction effect of sobetirome with bleomycin is rather profound: comprising 53.5% of the total differentially affected genes and the majority of bleomycin’s main effect features (**Fig.2C**), (**Table S1**).

**Figure 2.**
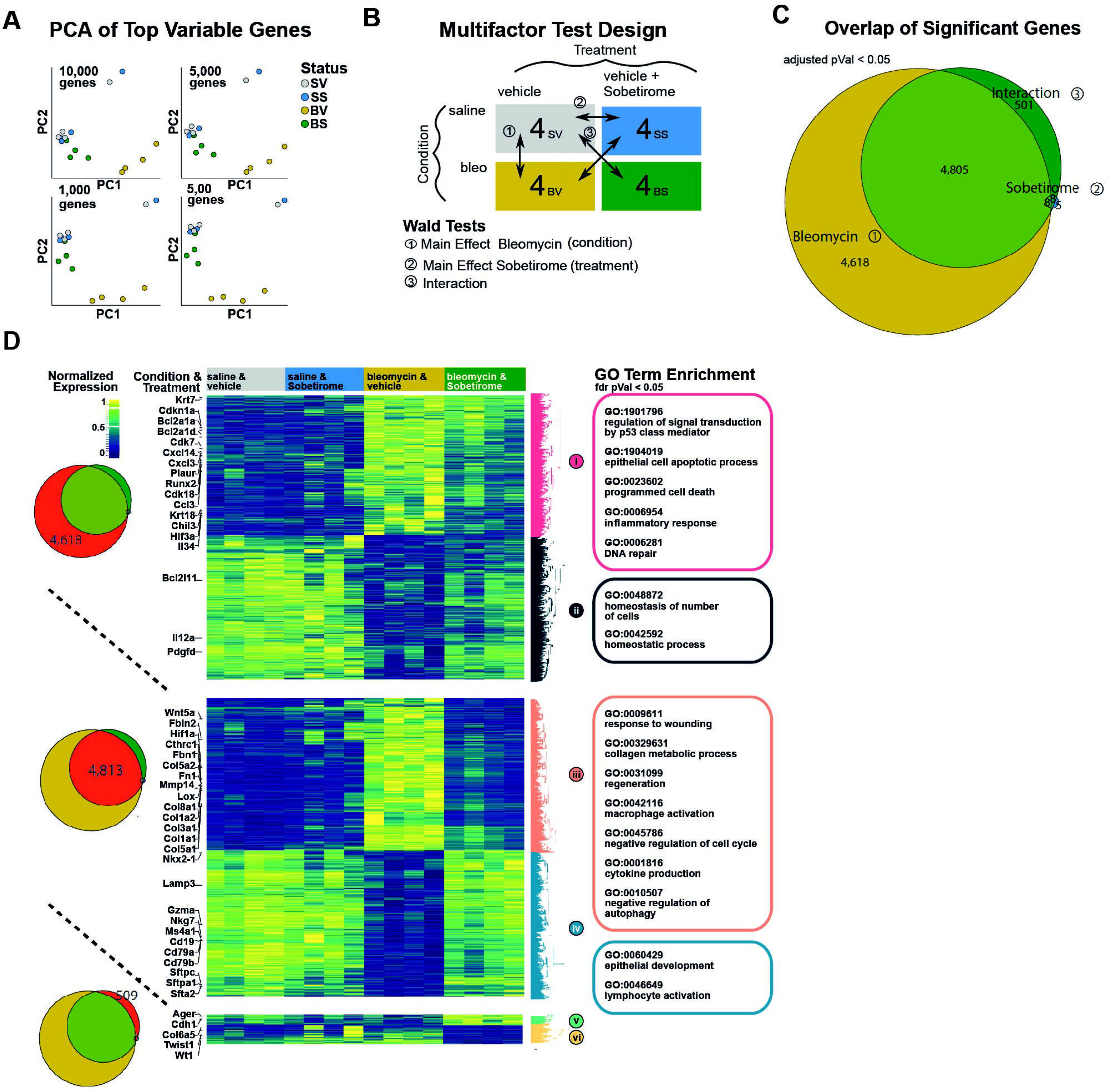
Transcriptional analysis demonstrates that sobetirome exerts minimal effects on healthy lungs but reverses fibrosis-associated gene signatures in mice following bleomycin exposure. **(A)** PCA of the top n genes ranked by variance. (**B)** Diagram of experimental design for multifactor testing. **(C)** Euler diagram of significant genes from each test in multifactor analysis. **(D)** Heatmap of significant gene sets from respective regions of the Euler diagram. Gene expression was subject to variance-stabilizing transformation before being min-max normalized across samples. Genes from each set were hierarchically clustered and segmented into two clusters. Significantly enriched GO terms from each cluster were selected for presentation.

We further dissected the molecular contributions of these gene sets and their relative expression changes across conditions. Independent of sobetirome treatment, bleomycin exposure induced a robust transcriptional response characterized by an increase in genes associated with apoptosis (*Bcl2a1a, Bcl2a1d*), inflammation (*Ccl3, Cxcl14, Il34*), and DNA damage/repair (*Cdkn1a, Plaur*), accompanied by a coordinated decrease in genes linked to homeostatic biological processes (**Fig. 2D, upper)**. Notably, analysis of the 4,813 genes shared between the bleomycin main-effect and interaction-effect gene sets revealed a striking reversal of these bleomycin-driven transcriptional changes following sobetirome treatment **(Fig. 2D, middle)**. Specifically, sobetirome markedly attenuated the expression of canonical fibrotic markers (*Col1a1, Col1a2, Col3a1, Col5a1, Fn1, Fbn1, Cthrc1, Lox, Mmp14*) while restoring expression of key epithelial and immune cell–associated genes, including alveolar type II (ATII) cell markers (*Sftpc, Sftpa1, Sfta2, Lamp3*), components of fibroblast growth factor receptor signaling (*Fgf10, Fgfr3*), B cell–associated genes (*Cd19, Cd79b*), and cytotoxic lymphocyte markers (*Nkg7, Gzma*) **(Fig. 2D, middle)**. In addition to these protective interactions with bleomycin-induced injury, sobetirome exerted significant interaction effects independent of bleomycin’s main effect. These included a sobetirome-driven enrichment of epithelial identity genes (*Ager, Cdh1*) and a concomitant reduction in select fibrosis-associated markers (*Col6a5, Twist1, Wt1*) **(Fig. 2D, lower)**. Thus, beyond mitigating injury-induced transcriptional remodeling, sobetirome independently promotes epithelial features while suppressing specific profibrotic programs.

In summary, our findings demonstrate that sobetirome exerts minimal transcriptional effects in uninjured lungs yet potently reverses fibrosis-associated gene expression in the context of bleomycin-induced injury. Moreover, sobetirome restores epithelial and immune cell signatures and independently enhances epithelial identity while reducing selected fibrotic markers. Together, these results underscore the safety of sobetirome in healthy lung tissue and highlight its robust antifibrotic and epithelial-protective activity following lung injury.

### Sobetirome Mitigates Bleomycin-Induced Alveolar Epithelial Cell Apoptosis and Mitochondrial Dysfunction

To delineate the mechanisms underlying the antifibrotic effects of sobetirome, we next investigated its impact on alveolar epithelial cell apoptosis and mitochondrial function. Building on our previous findings that thyroid hormone T3 reduces epithelial apoptosis in bleomycin-induced pulmonary fibrosis [18], we assessed whether sobetirome exerts similar protective effects.

Our study revealed that sobetirome treatment markedly reduces the proportion of TUNEL-positive cells in the lungs of bleomycin-challenged mice compared with bleomycin alone, an effect also confirmed by decreased levels of cleaved caspase-3, indicating attenuation of epithelial apoptosis **(Fig. S2A-E)**. Additionally, co-staining for pro–surfactant protein C (pro-SPC) and TUNEL revealed a significant increase in apoptotic ATII cells following bleomycin exposure, which was effectively abrogated by sobetirome treatment **(Fig. 3A, B).**

**Figure 3.**
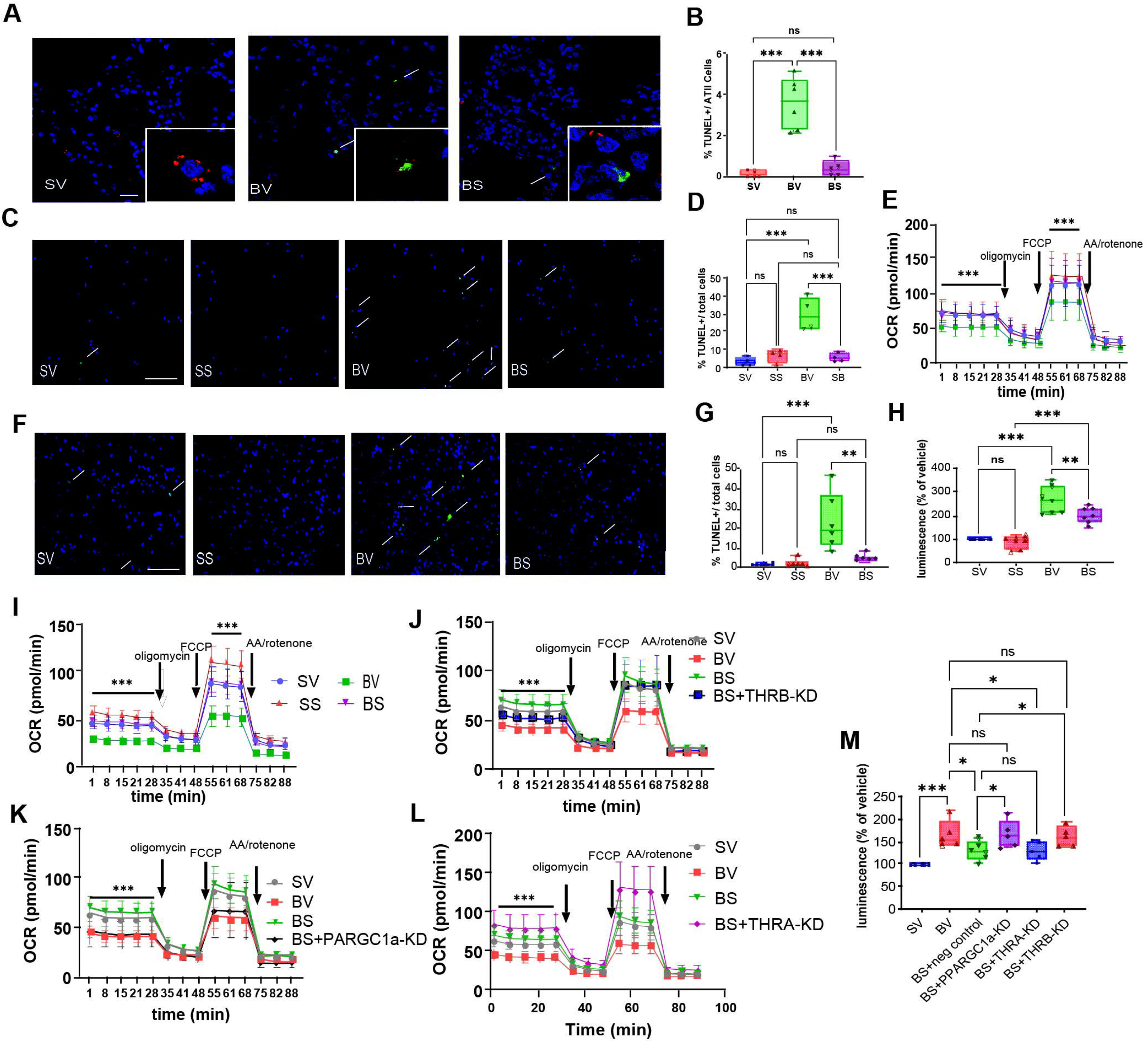
Sobetirome rescues epithelial cells from bleomycin-induced apoptosis and mitochondrial dysfunction. **(A)** Photomicrographs of mouse lungs treated with vehicle and saline (SV), vehicle and bleomycin (1,5U/kg, BV), and sobetirome and bleomycin (BS; 300 mg/kg every other day) stained for pro-SPC (red), TUNEL (green), and DAPI (blue). Arrows indicate double-positive cells. Insets show (SV) a pro-SPC positive/TUNEL negative cell; (BV) a pro-SPC positive/TUNEL positive cell; and (SB) a pro-SPC negative/TUNEL positive cell. **(B)** Double-positive cells were counted in a blinded fashion. **(C-E)** Alveolar type II cells were isolated from C57/Bl6 mice, treated with bleomycin (15mU/ml for 6 h) followed by sobetirome (100 µg/ml) for 18 h, and apoptosis was assessed via TUNEL staining. **(C, D)** Representative images and quantification analysis for ATII cells treated with vehicle and saline (SV), vehicle and bleomycin (15 mU/ml, BV), sobetirome and saline (SS), and sobetirome and bleomycin (BS). Arrows indicate double-positive cells. **(E)** Oxygen consumption rate for isolated ATII cells as measured by the Seahorse device. **(F-M)** Small airway epithelial cells were treated with bleomycin (15mU/ml for 6 h) followed by sobetirome (100 µg/ml) for 18 h and apoptosis was assessed via **(F, G)** representative images for cells treated with vehicle and saline (SV), vehicle and bleomycin (15 mU/ml, BV) sobetirome and saline (SS) and sobetirome and bleomycin (BS). Arrows indicate double-positive cells for TUNEL staining. **(H)** Caspase 3/7 activity as measured by luminescence. **(I)** Oxygen consumption rate for SAEC as measured by the Seahorse device. **(J-L)** SAECs were transfected with negative control siRNA or siRNA targeted to *PPARGC1α*, *THRA*, or *THRB* for 48 h. Negative controls (SV, BV) and respective siRNA-transfected cells were harvested after treatment with bleomycin (15 mU/ml for 6 h) followed by sobetirome (100 µg/ml) for 18 h. **(J, K, L)** OCR measurements in cells treated with bleomycin and sobetirome and transfected with siRNA for (**J**)THRB, **(K)** PPARGC1α, **(L)** THRA. **(M)** Apoptosis using caspase 3/7 luminescence. One-way ANOVA was performed for B, D, G, H, and M. Two-way ANOVA followed by the Holm-Sidak multiple comparison test was performed for E, I, J, K, and L (n=6 and 7, respectively). *=p<0.05, **=p<0.01, ***=p<0.001, ***=p<0.001, scale bars show (A) 20 µm and (C) 100µm, respectively.

To validate these observations at the cellular level, we assessed the effects of sobetirome in freshly isolated murine ATII cells **(Fig S3A-C).** Sobetirome significantly mitigated bleomycin-induced apoptosis and mitochondrial dysfunction in these cells, as evidenced by a reduction in TUNEL positivity and restoration of mitochondrial respiration, measured by oxygen consumption rate (OCR) **(Fig. 3C–E and Fig. S4A-F).**

Extending these findings to human cells, we evaluated the impact of sobetirome on human small airway epithelial cells (SAECs) exposed to bleomycin. Sobetirome partially rescued SAECs from bleomycin-induced apoptosis, as indicated by decreased TUNEL staining and reduced caspase-3/7 activity **(Fig. 3F–H)**. Consistent with these anti-apoptotic effects, sobetirome also restored mitochondrial function in bleomycin-treated SAECs **(Fig. 3I and Fig. S4G-L)**.

Finally, to define the molecular mediators of sobetirome in these epithelial-protective effects, we selectively knocked down *THRA*, *THRB*, and *PPARGC1A* in SAECs (**Fig. S5A-C)**. Loss-of-function analyses revealed that sobetirome-mediated reductions in apoptosis and restoration of mitochondrial function were dependent on THRB and PPARGC1A, but not THRA **(Fig. 3J–M).**

In summary, these data demonstrate that sobetirome robustly attenuates bleomycin-induced epithelial apoptosis and mitochondrial dysfunction through a THRB– PPARGC1A–dependent mechanism, providing mechanistic insight into its epithelial-protective and antifibrotic actions.

### Sobetirome Enhances Fibroblast Apoptosis and Mitochondrial Function in Lung Fibrosis

Given the overall reduction in apoptosis observed in lung tissues of bleomycin-challenged mice treated with sobetirome, we next examined whether this effect extended specifically to myofibroblasts. To this end, lung sections from bleomycin-exposed mice with or without sobetirome treatment were co-stained for α-smooth muscle actin (αSMA) and TUNEL. This analysis revealed a significant increase in αSMA/TUNEL double-positive cells in sobetirome-treated lungs, indicating enhanced myofibroblast apoptosis following bleomycin injury **(Fig. 4A).**

**Figure 4.**
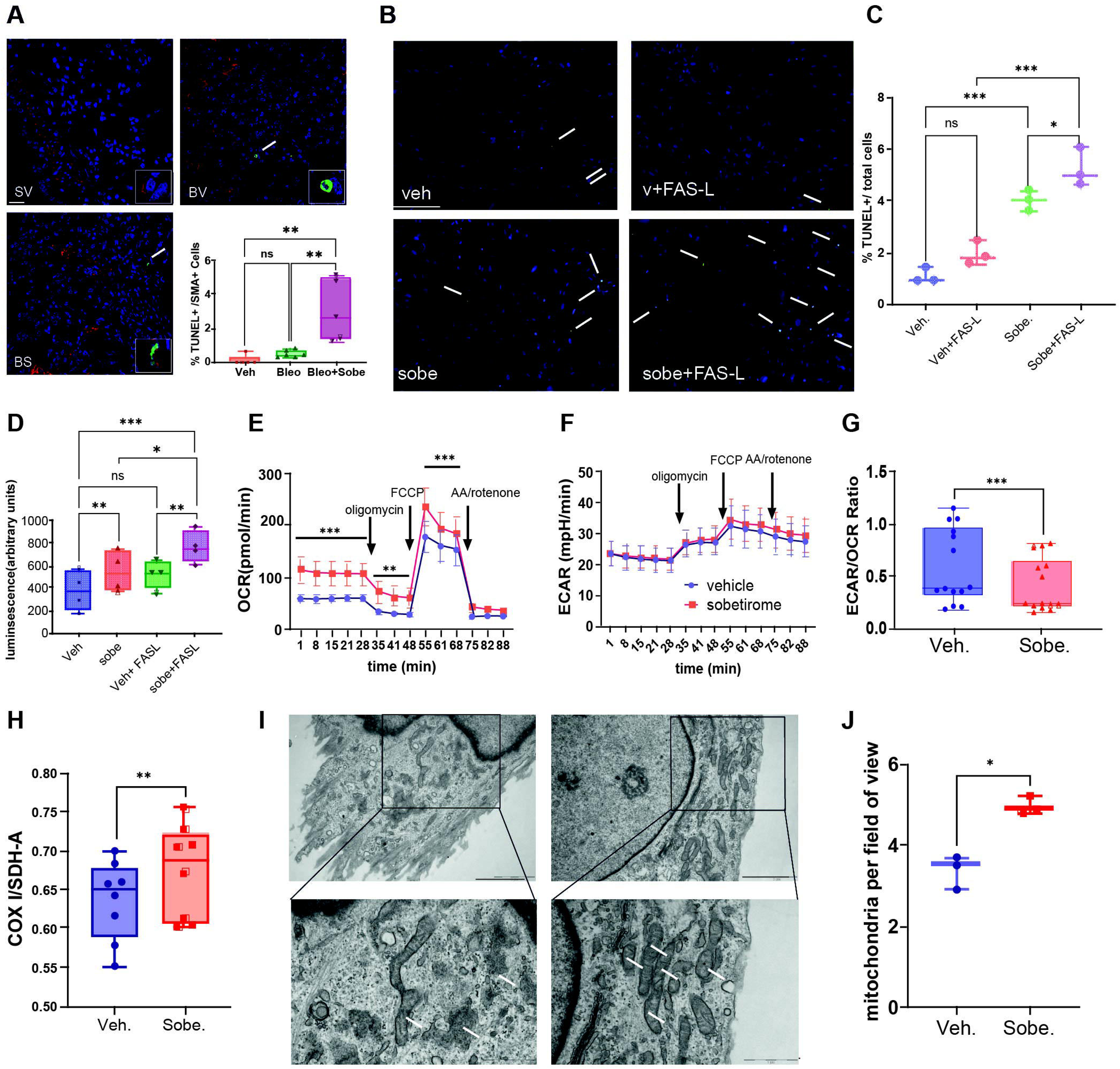
Sobetirome promotes fibroblast apoptosis and ameliorates mitochondrial dysfunction. **(A)** Photomicrographs of mouse lungs treated with vehicle and saline (SV), vehicle and bleomycin (1,5U/kg, BV), and sobetirome and bleomycin (BS; 300 mg/kg every other day) stained for aSMA (red), TUNEL (green), and DAPI (blue). Arrows indicate double-positive cells. Insets show (SV) an αSMA positive/TUNEL negative cell; (BV) an αSMA negative/TUNEL positive cell; and (SB) an αSMA positive /TUNEL positive cell. Double-positive cells were counted in a blinded fashion. **(B-D)** IPF fibroblasts were treated for 7 days with vehicle (veh) or sobetirome (100 ng/ml, sobe.), and vehicle and FAS-ligand (300 ng/ml, FAS-L) were added 24 hours before apoptosis was assessed via **(B** and **C)** representative images and quantifications for TUNEL staining; arrows indicate double-positive cells. **(D)** Caspase 3/7 activity as measured by luminescence. **(E)** Oxygen consumption rate (OCR) and **(E)** Extracellular acidification rate (ECAR) for IPF fibroblasts as measured by the Seahorse device. **(G)** Ratio of ECAR/OCR of IPF fibroblasts with and without sobetirome. **(H)** Mitobiogenesis was assessed as a ratio of COXI to SDHA protein. **(I)** Representative transmission electron microscopy (TEM) images (n = 21-25, respectively). **(J)** Number of mitochondria in TEM images. One-way ANOVA was performed for B, D, F, I, and H. Two-way ANOVA followed by the Holm-Sidak multiple comparison test was performed for E (n=6). Student’s t-test was performed for I and K. *=p<0.05, **=p<0.01, ***=p<0.001. Scale bars show (A) 20 µm, (C) 100µm, and (J) 2 µm (500nm in inserts), respectively.

Fibroblasts derived from IPF lungs are well known to acquire resistance to apoptosis during fibrogenesis and to rely on anti-apoptotic pathways for survival [28, 29]. To determine whether sobetirome directly modulates apoptosis in this apoptosis-resistant cell population, we evaluated its effects on primary fibroblasts isolated from IPF lungs. Sobetirome treatment significantly increased apoptosis in IPF fibroblasts both in the presence and absence of Fas ligand stimulation, as demonstrated by increased TUNEL positivity and elevated caspase-3/7 activity **(Fig. 4B-D)**.

We further aimed to assess the effects of sobetirome on mitochondrial function in IPF fibroblasts and identified a significant improvement in mitochondrial function in these cells after sobetirome treatment, as indicated by OCR as well as all other key mitochondrial parameters (basal respiration, proton leak, peak respiration, spare respiratory capacity, ATP-linked respiration, and non-mitochondrial oxygen consumption) (**Fig. 4E** and **Fig. S6A-F)**. The extracellular acidification rate (ECAR) is an indicator of glycolysis, and it has been previously reported that the ECAR/OCR ratio is elevated in IPF fibroblasts and TGF-β-treated normal human lung fibroblasts (NHLFs) [13]. We observed that although sobetirome treatment did not alter the level of ECAR in IPF fibroblasts **(Fig. 4F),** the ECAR/OCR ratio was significantly reduced after sobetirome treatment (P-value ≤ 0.0001) **(Fig. 4G)**. This suggests that sobetirome induced a metabolic shift toward enhanced mitochondrial oxidative phosphorylation in this setting. We further confirmed the effects of sobetirome on mitochondrial homeostasis in IPF fibroblasts by showing an increase in mitochondrial biogenesis **(Fig. 4H)** as well as an increase in the number of mitochondria in these cells after sobetirome treatment using transmission electron microscopy (TEM) **(Fig. 4I and J)**.

These findings reveal that sobetirome promotes apoptosis in myofibroblasts, improves mitochondrial function, and stimulates mitochondrial biogenesis in lung fibrosis.

### Sobetirome Induces Apoptosis and Enhances Mitochondrial Function in IPF Fibroblasts via AMPK Phosphorylation, THRB, and PPARGC1α

Although enhanced mitochondrial function is commonly associated with decreased apoptosis, recent studies suggest that AMP-activated protein kinase (AMPK)-activating metabolic modulators, such as metformin and AICAR, can uncouple these processes by simultaneously improving mitochondrial function and inducing apoptosis in IPF fibroblasts [14]. Consistent with these findings, thyroid hormone signaling has also been shown to promote AMPK phosphorylation [30, 31], suggesting a potential convergence on shared metabolic pathways regulating fibroblast fate and mitochondrial function.

To further delineate the downstream pathways mediating the observed effects of sobetirome in IPF fibroblasts, we examined its impact on AMPK activation and PPARGC1α expression. Our study confirmed that sobetirome treatment significantly increases AMPK phosphorylation and upregulates PPARGC1α expression in fibroblasts derived from IPF patients **(Fig. 5A–C).** Notably, the magnitude and pattern of these effects closely resembled those induced by AICAR, suggesting engagement of a shared AMPK-dependent metabolic pathway.

**Figure 5.**
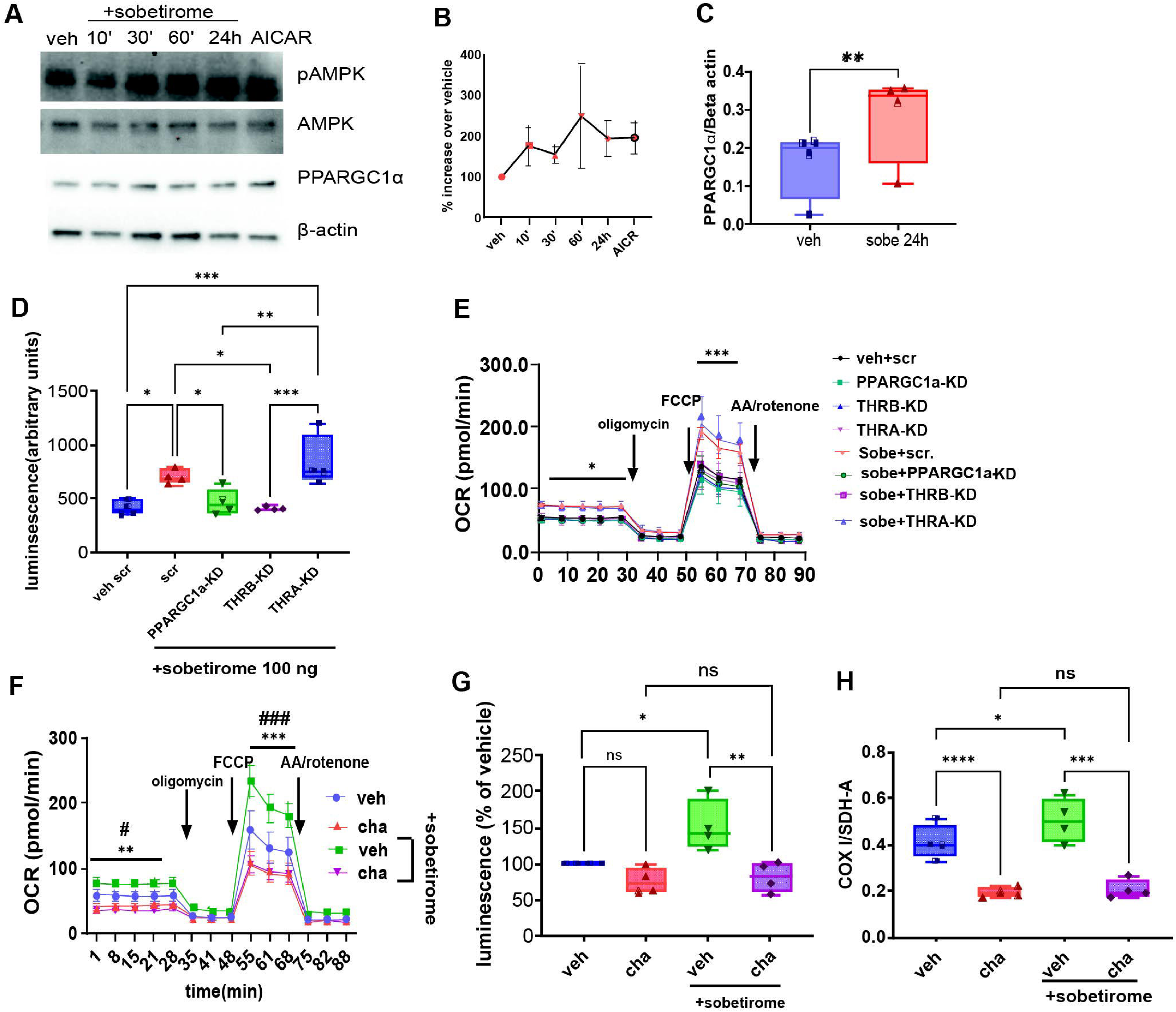
The effects of sobetirome on IPF-fibroblasts are mediated via mitobiogenesis, thyroid hormone receptor beta, and PPARGC1α. (**A**) Representative western blot showing phosphorylated AMP-activated protein kinase (pAMPK), total AMPK, PPARGC1α, and beta-actin in sobetirome (100 ng/ml, sobe.) treated IPF fibroblasts at different time points and AICAR (1mM) after 24 h. **(B)** Ratio of phosphorylated AMPK to AMPK. **(C)** Relative levels of PPARGC1α after 24 h of sobetirome. **(D-E**) Fibroblasts were transfected twice with negative control siRNA (neg control) or siRNA targeted to *PPARGC1α*, *THRA,* or *THRB* and were assessed after 7 days. **(D)** Apoptosis was measured in IPF fibroblasts transfected with siRNA to respective targets or negative control siRNA in the presence of sobetirome. **(E)** Oxygen consumption rate (OCR) was measured in IPF fibroblasts transfected with siRNA to respective targets or negative control siRNA in the presence of sobetirome. **(F-H)** IPF fibroblasts were treated with chloramphenicol (cha, 300 µM) for 7 days, both in the presence and absence of sobetirome, and **(F)** Oxygen consumption rate (OCR), **(G)** apoptosis, and **(H)** mitobiogenesis were assessed. One-way ANOVA was performed for D, G, and H. Two-way ANOVA followed by the Holm-Sidak multiple comparison test was performed for E and F (n=5 and 6, respectively). Student’s t-test was performed for C.*=p<0.05, **=p<0.01, ***=p<0.001, =p<0,05 veh vs cha, =p<0.001 veh vs cha.

Next, we sought to characterize the receptor and downstream effector signaling mechanisms mediating the cellular responses to sobetirome in IPF fibroblasts. To guide our mechanistic studies, we reanalyzed our previously published single-cell RNA sequencing data from IPF lung tissues [32]. This transcriptomic analysis revealed increased expression of *THRB* in myofibroblasts and alveolar fibroblasts from IPF lungs compared with controls **(Fig. S7A).** We confirmed these observations by in situ hybridization for THRB combined with αSMA co-staining in IPF lung sections **(Fig. S7B).** Consequently, we targeted siRNA-mediated knockdown of *THRB*, *THRA*, and *PPARGC1A* in IPF fibroblasts **(Fig. S8A-C).** This study demonstrates that the pro-apoptotic and mitochondrial effects of sobetirome are strictly dependent on *THRB* and *PPARGC1A* signaling, whereas *THRA* is non-essential for these cellular responses **(Fig. 5D, E)**.

We next aimed to determine whether the effects of sobetirome on mitochondrial biogenesis in IPF fibroblasts (as shown in Figure 4H-J) are necessary for its observed effects on mitochondrial function and apoptosis. To address this question, we used chloramphenicol, a well-established inhibitor of mitochondrial protein synthesis widely used to suppress mitochondrial biogenesis. [33, 34]. Seahorse analysis revealed that chloramphenicol significantly reduces OCR both in the presence and absence of sobetirome, with no significant differences observed between chloramphenicol-treated groups at any time point **(Fig. 5F).** Interestingly, chloramphenicol at the used concentration did not affect apoptotic indices but completely abolished the sobetirome-induced increase in apoptosis in these cells **(Fig. 5G).** Importantly, consistent with its known mechanism of action, chloramphenicol markedly suppressed mitochondrial biogenesis irrespective of sobetirome treatment **(Fig. 5H).** Together, these findings indicate that mitochondrial biogenesis is essential for sobetirome-mediated enhancement of mitochondrial function and induction of apoptosis in IPF fibroblasts.

In summary, sobetirome promotes apoptosis and improves mitochondrial function in IPF fibroblasts through a THRB-dependent signaling axis involving AMPK phosphorylation and PPARGC1α-driven mitochondrial biogenesis.

### PPARGC1α Expression in ATII and Col1a1-Positive Cells is Essential for the Antifibrotic Effects of Sobetirome

Collectively, our findings demonstrate robust antifibrotic effects of sobetirome across multiple lung cell populations, including IPF fibroblasts, ATII epithelial cells, and SAEC. We have previously shown that Ppargc1α is a critical mediator of thyroid hormone signaling in the lung and that global knockdown of Ppargc1α exacerbates bleomycin-induced pulmonary fibrosis [18]. Building on these observations, we demonstrate here that Ppargc1α is also a key downstream effector of sobetirome in both ATII cells and IPF fibroblasts.

To further define the cell-type–specific role of Ppargc1α in mediating the antifibrotic actions of sobetirome, we performed co-immunostaining for Ppargc1α in combination with either pro-SPC or collagen I in lung tissues from bleomycin-challenged mice. We discovered that sobetirome treatment is associated with a trend toward increased Ppargc1α expression and enhanced nuclear localization in both ATII cells and Col1a1-positive fibroblasts **(Fig. S9),** supporting activation of Ppargc1α-dependent transcriptional programs in these populations during fibrosis resolution.

Next, we generated mice harboring floxed *Ppargc1α* alleles crossed with either *Col1a1-Cre/ERT2* or *Sftpc-Cre/ERT2* drivers to enable cell-type–specific deletion of Ppargc1α and directly assess its requirement for sobetirome-mediated antifibrotic effects in the bleomycin mice model **(Fig. 6A).** Notably, cell-specific deletion of *Ppargc1α* alone did not exacerbate fibrosis, as assessed by weight loss, lung mechanics, hydroxyproline content, and histopathological analyses **(Fig. 6B–H).** In contrast, deletion of *Ppargc1α* in either *Col1a1*- or *Sftpc*-positive cells was sufficient to abolish the antifibrotic effects of sobetirome, as evidenced by greater weight loss, worsened histological fibrosis, increased collagen accumulation, and reduced lung function parameters following bleomycin challenge **(Fig. 6B–H).**

**Figure 6.**
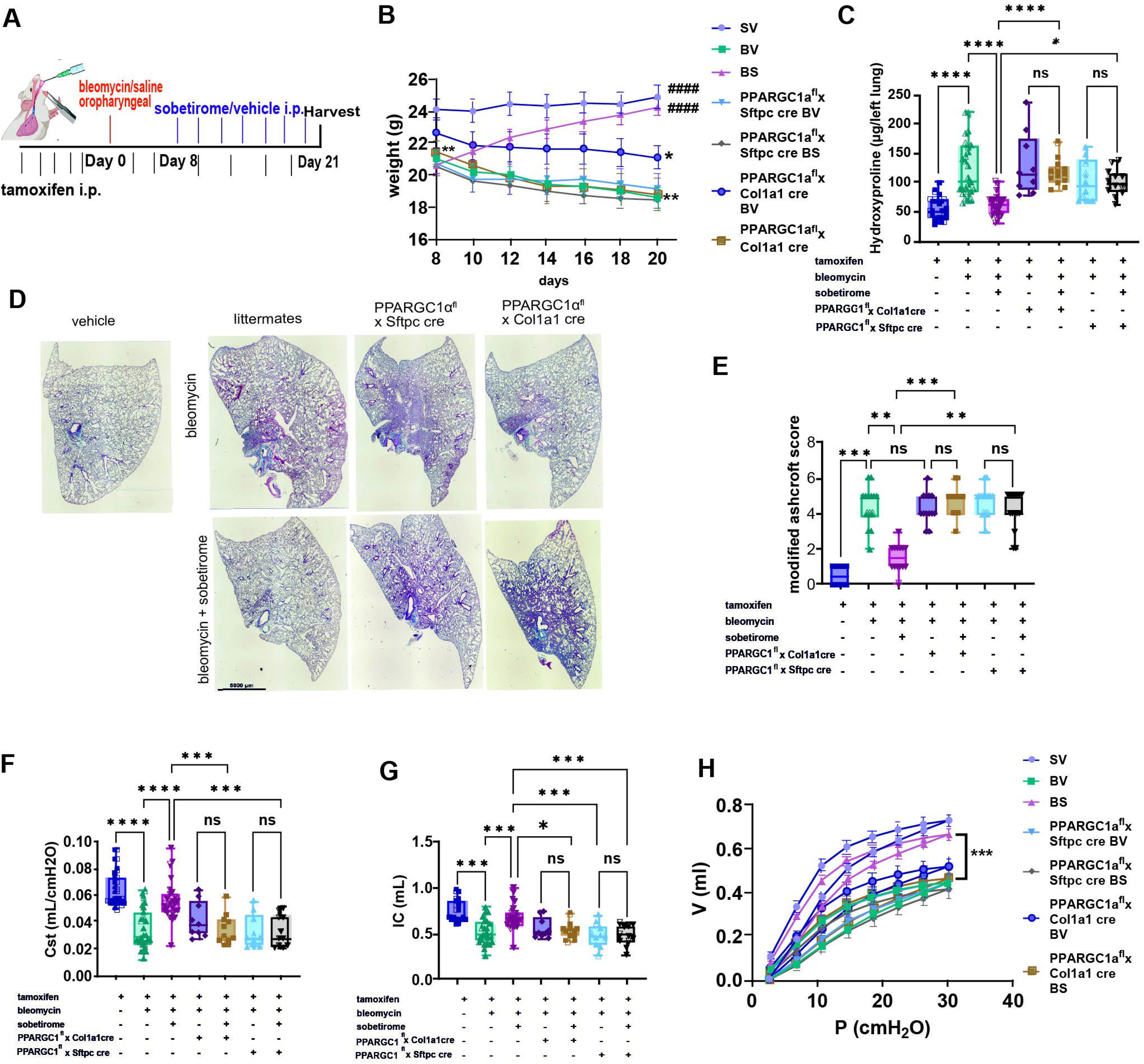
*Ppargc1α* expression in both ATII and *Col1a1*-positive cells is required for the antifibrotic effects of sobetirome in mice. **(A)** Experimental Design. **(B)** Total Body Weight was measured every other day, starting at the beginning of sobetirome/vehicle treatment, and the last observation was carried forward for deceased mice. **(C)** Hydroxyproline levels, as a measurement for lung collagen, were assayed from lung lysates. **(D-E)** Histological assessments of the mouse lung. Whole lobes of mice of the indicated groups were stained with Masson’s trichrome. The upper row shows bleomycin and vehicle (BV), and the lower row shows bleomycin and sobetirome (BS) treated mice, while the insert on the right shows vehicle (SV) only. Photomicrographs of trichrome-stained sections were taken and graded in a blinded fashion. **(F-H)** Mice lung function assessments using FlexiVent System: **(F)** Static compliance, **(G)** total lung capacity, and **(H)** Pressure-Volume curves were assessed. One-way ANOVA was performed for C, F, and G, and the Kruskal-Wallis test for E. Two-way ANOVA followed by the Holm-Sidak multiple comparison test was performed for B and H (n=4-14). Scale bar: 5mm *=p<0.05, **=p<0.01, ***=p<0.001, in E**=p<0.01 veh vs respective conditions and ##=p<0.01 bleomycin only vs respective condition.

Together, these results establish Ppargc1α as an essential, cell-intrinsic mediator of sobetirome in pulmonary fibrosis and provide a mechanistic bridge between its metabolic effects at the cellular level and its antifibrotic activity *in vivo*.

### The inhalational administration of sobetirome effectively ameliorates bleomycin-induced pulmonary fibrosis

Our study showed that IP administration of sobetirome attenuates lung fibrosis and does not result in significant weight loss, hepatotoxicity, or other clinical features of thyrotoxicosis. Nevertheless, given the potential for systemic side effects with chronic treatment, we next evaluated the feasibility and therapeutic efficacy of a localized, non-systemic delivery approach. To this end, we assessed the effects of inhalational delivery of sobetirome via daily aerosolized administration following bleomycin challenge **(Fig. 7A).**

**Figure 7.**
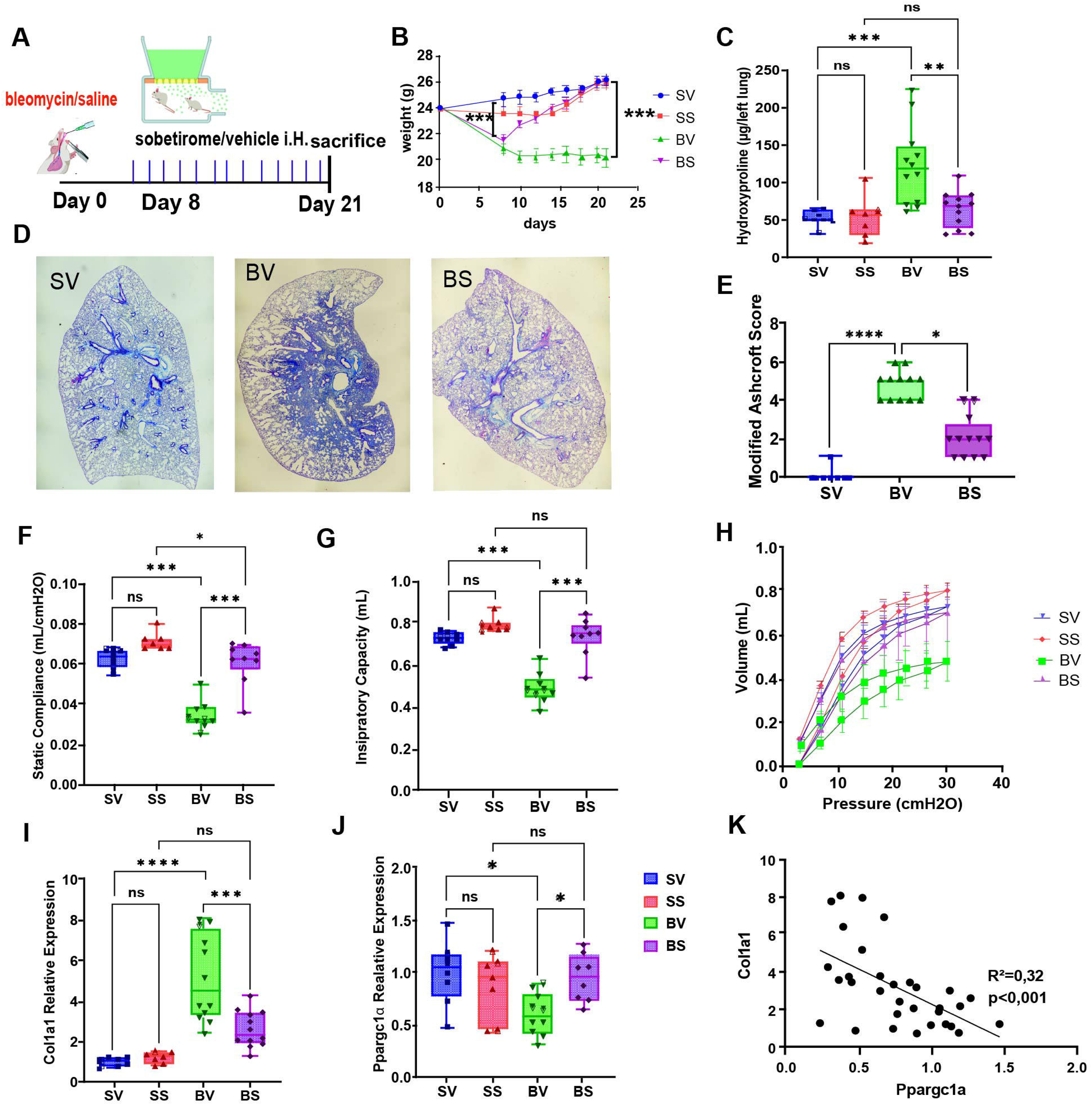
Inhalational administration of sobetirome ameliorates pulmonary fibrosis in a mouse model. **(A)** Experimental Design. **(B)** Total body weight was measured every other day, starting at the beginning of sobetirome/vehicle treatment, and the last observation was carried forward for deceased mice. **(C)** Hydroxyproline levels, as a measurement for lung collagen, were assayed from lung lysates (sobetirome and saline, SS). **(D-E)** Lung Histological Assessments: Whole lobes of mice treated with vehicle and saline (SV), vehicle and bleomycin (1,5U/kg, BV), and sobetirome and bleomycin (BS; 300 mg/kg every other day) were stained with Masson’s trichrome. Scale bars indicate 1 mm. Photomicrographs of trichrome-stained sections were taken and graded in a blinded fashion. **(F-H)** Lung function measurements using the FlexiVent system: **(F)** Static compliance, **(G)** Inspiratory capacity, and **(H)** Pressure-Volume curves. (I-K) Gene expression measurements by qPCR of lung tissue for **(I)** *Col1a1* and **(J)** *Ppargc1α*. **(K)** Linear regression analysis showing the relative expression of *Col1a1* and *Ppargc1α*. One-way ANOVA was performed for C, F, G, I, J, and the Kruskal-Wallis test for E. Two-way ANOVA followed by the Holm-Sidak multiple comparison test was performed for B and H (n=12-23 and 7-8, respectively). scale bar: 1mm *=p<0.05, **=p<0.01, ***=p<0.00.

Inhaled sobetirome treatment promoted more rapid and complete recovery of body weight compared with IP administration **(Fig. 7B)** and resulted in significant improvements in lung histopathology and reductions in hydroxyproline content in bleomycin-injured mice **(Fig. 7C–E).** Additionally, lung functional assessments further demonstrated marked improvements in static compliance, inspiratory capacity, and pressure–volume relationships after sobetirome treatment, compared to vehicle-treated animals **(Fig. 7F–H).** Furthermore, sobetirome treatment reversed the bleomycin-induced increase in *Col1a1* mRNA expression and the increase in *Ppargc1α* gene expression after bleomycin (**Fig. 7I** and **J**). Additionally, we found a significant inverse correlation between *Ppargc1α* and *Col1a1* expression levels (R² = 0.32, p < 0.001) in these mice lungs **(Fig. 7K)**. This finding further supports a mechanistic link between Ppargc1α activation and suppression of fibrotic gene expression.

Collectively, these findings establish inhalational delivery of sobetirome as a targeted, efficacious, and potentially safer therapeutic strategy for the treatment of pulmonary fibrosis.

### Sobetirome exerts antifibrotic effects in human lung fibrosis using an *ex vivo* precision-cut lung slice model

Building on the robust antifibrotic effects of sobetirome observed in murine models and *in vitro* cellular systems, we next evaluated its therapeutic potential in an *ex vivo* human precision-cut lung slice (hPCLS) model. We first assessed the effects of sobetirome on hPCLS obtained from healthy donor lungs cultured in the presence of either a profibrotic cocktail (FC; TGF-β (Transforming growth factor-beta), PDGF-α (Platelet-derived growth factor subunit A), TNF-α (Tumor Necrosis Factor-alpha), and LPA (lysophosphatidic acid)) or a control cocktail, as previously described **(Fig. 8A)** [35, 36]. Exposure to the FC resulted in a marked induction of *COL1A1* mRNA expression and an increase in secretion of Pro-COL1A1 protein, both of which were significantly attenuated by sobetirome treatment **(Fig. 8B** and **C**). Consistent with these findings, sobetirome also led to a pronounced reduction in collagen deposition, as demonstrated by Masson’s trichrome staining in these slices **(Fig. 8D, E)**.

**Figure 8.**
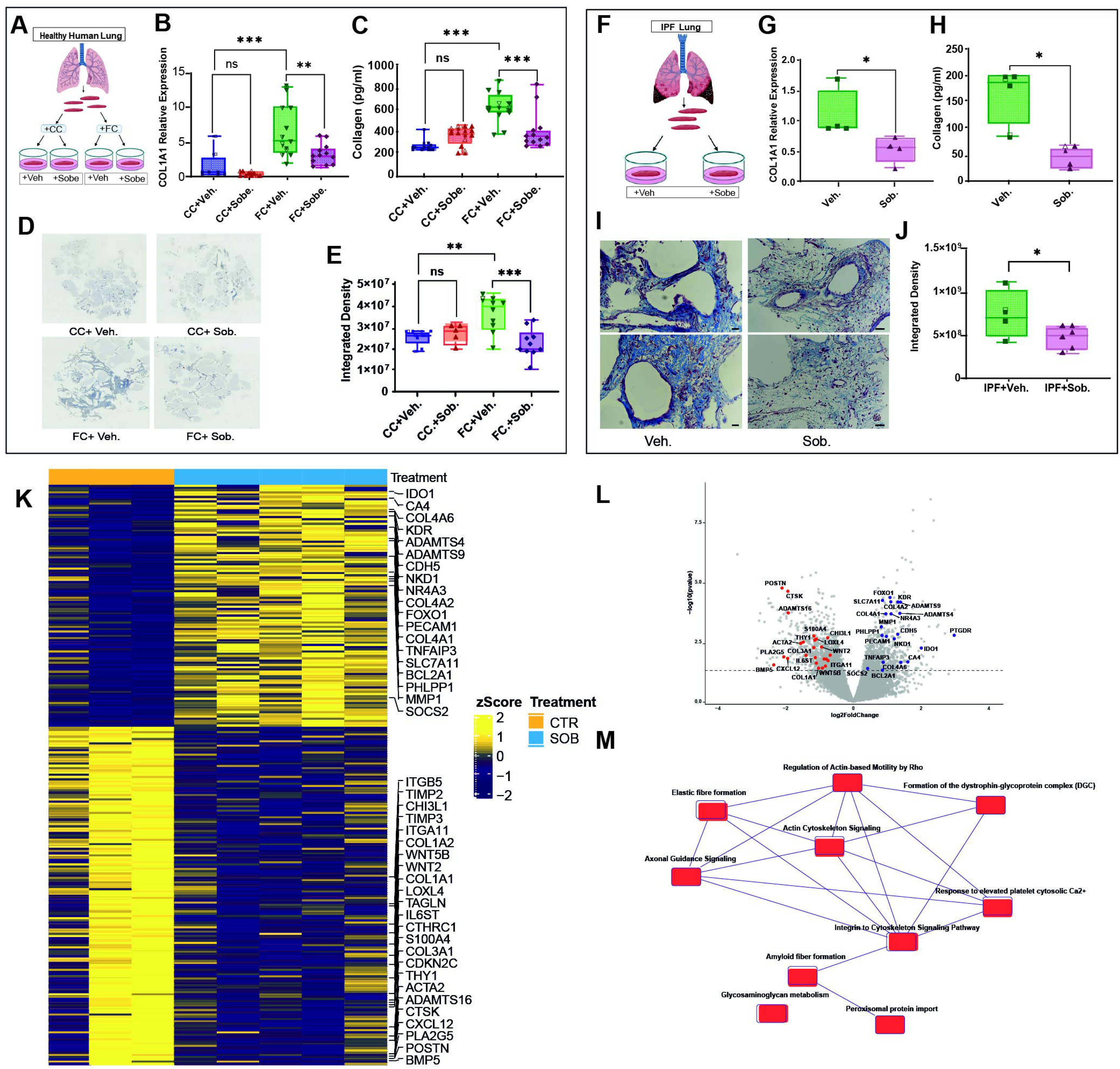
Sobetirome attenuates fibrotic responses in human precision-cut lung slices (hPCLS). **(A-E)** hPCLS obtained from healthy donor lungs: **(A)** Schematic of experimental design: hPCLS from healthy donor lungs were cultured with either a profibrotic cocktail (FC; TGF-β, PDGF-α, TNF-α, and LPA) or control conditions. **(B)** *COL1A1* mRNA expression; **(C)** Pro-COL1A1 secretory protein measurement by ELISA. **(D–E)** Representative images and quantification of Masson’s trichrome staining demonstrate increased collagen deposition following FC treatment, which is substantially attenuated by sobetirome. **(F-J)** hPCLS obtained from IPF lungs: **(F)** Experimental design. **(G)** *COL1A1* gene expression; **(H)** secreted Pro-COL1A1 protein levels. **(I–J)** Masson’s trichrome staining and corresponding quantification confirm reduced collagen accumulation in sobetirome-treated IPF hPCLS. **(K–M)** Bulk RNA sequencing of IPF hPCLS: **(K)** Heatmaps of significant Differentially expressed genes (DEGs) between control vs sobetirome-treated slices. **(L)** Volcano plot illustrating DEGs (x-axis: log₂ fold change; y-axis: −log₁₀ *P* value). **(M)** Canonical pathway overlap analysis using Ingenuity Pathway Analysis (IPA) demonstrates connections among downregulated fibrotic signaling pathways, indicating shared molecular components and coordinated network regulation in response to sobetirome. Pathway overlap is defined by the proportion of shared genes relative to total genes within each pathway, with statistical significance determined by a right-tailed Fisher’s exact test (*P* < 0.05).

To validate these findings in a more disease-relevant context, we examined the effects of sobetirome in hPCLS derived from patients with IPF **(Fig. 8F)**. After five days of treatment, sobetirome markedly reduced the expression of the *COL1A1* gene in these lung slices as well as the secretory pro-COL1A1 protein in their supernatants **(Fig. 8G** and **H**). We further confirmed these antifibrotic effects of sobetirome by reduction in collagen accumulation in these IPF lung slices measured by Masson’s trichrome staining **(Fig. 8-I** and **J)**.

To gain broader insight into the transcriptional programs modulated by sobetirome in human lung fibrosis, we performed bulk RNA sequencing on IPF hPCLS following sobetirome treatment. This analysis revealed a coordinated antifibrotic transcriptional response characterized by both induction and repression of key gene programs. Upregulated genes, including *MMP1*, *ADAMTS4*, and *ADAMTS9*, indicate enhanced extracellular matrix turnover, while increased expression of *COL4A1*, *COL4A2*, and *COL4A6* suggests restoration of basement membrane architecture. Elevated endothelial markers (*PECAM1*, *CDH5*, *KDR*) further point to improved vascular integrity, alongside increased expression of regulatory and immunomodulatory genes (*TNFAIP3*, *FOXO1*, *NR4A3*, *IDO1*). In contrast, downregulation of profibrotic and myofibroblast-associated genes, including *COL1A1*, *COL1A2*, *COL3A1*, *ACTA2*, *TAGLN*, *POSTN*, *CTHRC1*, *S100A4*, *CTSK,* and *WNT* pathway components (*WNT2*, *WNT5B*), as well as matrix regulators (*TIMP2*, *TIMP3*, *LOXL4*), reflects suppression of extracellular matrix deposition and fibrotic signaling **(Fig. 8K** and **L) (Table S2)**.

Deeper analysis using Ingenuity Pathway Analysis (IPA) revealed that sobetirome modulates multiple key pathways relevant to IPF, with a consistent pattern of suppression across pathways implicated in disease progression. These include integrin–cytoskeleton signaling, glycosaminoglycan metabolism, elastic fiber formation, actin cytoskeleton signaling, β-catenin–independent WNT signaling, oxidative stress–induced senescence, the senescence-associated secretory phenotype (SASP), and inhibition of matrix metalloproteases (**Fig. S9A)**. Canonical pathways overlap analysis further demonstrated that many of these pathways share common biological components, indicating coordinated regulation across fibrotic networks **(Fig. 8M)**. Additionally, integration of canonical pathways with upstream regulator analysis identified key signaling nodes, showing that sobetirome suppresses downstream signaling of EGF (Epidermal Growth Factor), TEAD (Transcriptional Enhanced Associated Domain), VEGFA (*Vascular Endothelial Growth Factor A), and TGF-β, including pathways linked to fibrogenesis and cancer-associated processes **(Fig. S9B)**.

Collectively, these transcriptional changes indicate a shift from active fibrosis toward matrix remodeling, reduced myofibroblast activation, resolution of inflammation, and restoration of tissue homeostasis in the IPF lung. These findings highlight the therapeutic potential of sobetirome and support its continued investigation as a candidate treatment for IPF.

## Discussion

In this study, we identified sobetirome, a selective thyroid hormone receptor beta (THRB) agonist, as a potential antifibrotic agent through comprehensive *in vivo*, *in vitro*, and *ex vivo* preclinical studies in both human and murine models. Our findings demonstrate that the antifibrotic effects of sobetirome are mediated via *THRB* and *PPARGC1α*, with the expression of *PPARGC1α* in both ATII cells and Col1a1-positive cells being essential. Notably, we showed for the first time that sobetirome enhances mitochondrial function, mitigates apoptosis resistance, and promotes mitochondrial biogenesis in IPF fibroblasts. Most significantly, we provided evidence of sobetirome-induced improvements in human end-stage IPF tissue, including profound transcriptomic alterations.

Sobetirome has been shown to exert beneficial effects in lipid metabolism and has accordingly been tested in human phase I studies without obvious harmful side effects [26, 37]. However, the antifibrotic effects of this compound have been understudied; In this study, we show that, similar to T3 [18], sobetirome exerts a broad range of protective effects via the *THRB*, *PPARGC1α*, and mitochondrial function axis. Notably, these findings are particularly compelling because both fibroblasts and epithelial cells are affected, cell types that are widely recognized as key contributors to the pathogenesis and progression of IPF [10, 11]. Therefore, a drug capable of both protecting the alveolar epithelium and restoring IPF fibroblasts to a more physiological state may hold significant promise for future therapeutic applications and clinical trials [12].

We have already shown the protective effects of T3 in ATII cells [18], and a recent publication further highlighted the beneficial effects of T3 on these cells at the single-cell level in bleomycin-induced fibrosis [38]. In this study, we demonstrate that sobetirome, similar to T3, protects type II alveolar epithelial cells *in vitro* and *in vivo*. Using siRNA, we show for the first time that these effects are mediated via *THRB* and *PPARGC1α* but not *THRA* and go hand in hand with improvements in mitochondrial function. Given the fact that PPARGC1A serves as the “master regulator of mitobiogenesis” and T3 has long been implicated in improvements in mitochondrial function, these findings highlight the therapeutic potential of selectively targeting this pathway. This mechanism is strongly supported by our observation of parallel transcriptomic or phenotypic alterations in both murine ATII and human small airway epithelial cells. Furthermore, preserving alveolar epithelial integrity represents a highly promising therapeutic strategy, given that these cells exert diverse homeostatic functions, including the paracrine suppression of fibroblasts [39], and the regeneration of damaged epithelial cells [40]. Notably, while currently approved antifibrotic therapies primarily target fibroblast activation, some secondary epithelial-protective effects have been reported for nintedanib [41].

Our finding that sobetirome treatment significantly increased the apoptosis of aSMA-positive cells in sobetirome-treated fibrotic mouse lungs was somewhat unexpected. We first believed that this might be due to increased ATII cell viability and subsequent paracrine suppression of the fibroblasts [39, 42]. However, even in cell culture, the effect of sobetirome on apoptosis, both in the presence and absence of FAS-ligand, persisted. Intriguingly, this apoptotic response coincided with an increase in mitochondrial function within IPF fibroblasts, mirroring the metabolic enhancement observed in ATII cells. A similar metabolic and phenotypic shift has been documented with metformin and direct AMPK activators [14], a kinase that can increase *PPARGC1α* expression [43] and is, in turn, regulated by thyroid hormone [30]. Consistently, our data demonstrate that sobetirome alters both AMPK phosphorylation and PPARGC1A protein levels. Furthermore, the functional necessity of this axis was confirmed by targeted silencing, wherein knockdown of *THRB* or *PPARGC1A* but not *THRA* completely abrogated the effects of sobetirome.

While mitochondrial dysfunction is classically used as a hallmark of apoptosis, a growing body of evidence paradoxically demonstrates that restoring or enhancing mitochondrial bioenergetics can trigger apoptosis in activated, disease-associated fibroblasts [14, 44]. Cells with higher mitochondrial content, a feature mimicked by sobetirome-treated IPF fibroblasts, were shown to be more prone to apoptosis [50]. Another possible mechanism could be linked to the Warburg effect, a term describing the observation that some cells prefer glycolysis over oxidative phosphorylation, a feature that seems to be protective from apoptosis and can be reversed by metformin [45]. Importantly, this has previously been described for IPF fibroblasts [46] and coincides with our finding that sobetirome shifted this balance towards oxidative phosphorylation (decreased ECAR/OCR ratio). To prove that increased mitochondrial function mediates increased apoptosis susceptibility in response to sobetirome, we blocked mitobiogenesis using chloramphenicol. Importantly, this compound has been shown not to significantly affect glucose metabolism in human fibroblasts on its own, which is in contrast to the widely used alternative, ethidium bromide [47]. As expected, chloramphenicol reduced oxygen consumption rate, and there was no difference between chloramphenicol and sobetirome, showing that increased mitochondrial function is dependent on this process. More importantly, the effect of sobetirome on apoptosis was also abrogated in the presence of chloramphenicol.

Notably, the beneficial effects of *PPARGC1α* expression in IPF fibroblasts have been previously reported, whereas its silencing in normal human lung fibroblasts has been associated with a more fibrotic phenotype [48]. Interestingly, T3 stimulation did not result in a significant increase in *PPARGC1α* in these experiments. We assume that this discrepancy could be caused by the differing receptor specificity of T3 and sobetirome. In our hands, we found that while knockdown of *THRA* did not block the actions of sobetirome in IPF fibroblasts, it seemed to lead to a slight additional increase in oxygen consumption rate, a finding that, although likely caused by different mechanisms, had been previously observed in *THRA* knockout mice [49].

These observations led us to investigate the respective contribution of ATII cells and fibroblasts to the effects of sobetirome *in vivo*. As these were mediated via *PPARGC1α in vitro* in both cell types, we used *PPARGC1α^fl^* and *Sftpc-Cre/ERT2* as well as *Col1a1-Cre/ERT2* mice and induced excision of floxed alleles by a previously published protocol [50]. While Col1a1 is not exclusively expressed in fibroblasts and shows expression in smooth muscle cells and pericytes, it has recently been shown to label all fibroblast subpopulations in bleomycin-induced fibrosis [51]. In addition, pericytes have been shown to transdifferentiate towards myofibroblasts [52], and this might also be true for smooth muscle cells [53]. Furthermore, the widely used alternative marker for fibroblasts, *PDGFRα*, was not expressed in all subpopulations [51], and most importantly, is located on the same chromosome (5) as PPARGC1α, making the breeding of Cre-lox mice unfeasible. While total body knockdown of PPARGC1α increases bleomycin-induced hydroxyproline content, neither of the cell type-specific knockout mice showed any similar trend [18]. Notably, genetic ablation of *PPARGC1α* in either alveolar epithelial cells or fibroblasts completely abolished the antifibrotic effects of sobetirome, indicating that *PPARGC1α*-dependent signaling in both cell types is essential for its therapeutic activity. We did not observe evidence of residual protection in either knockout model, suggesting that the beneficial effects of sobetirome are largely dependent on intact *PPARGC1α* expression. Although we cannot formally exclude the possibility of a subtle effect that fell below the detection threshold, the inherent variability and regional heterogeneity of the bleomycin model make the identification of small differences particularly challenging [54]. Importantly, our findings do not support a biologically meaningful antifibrotic effect of sobetirome in the absence of *PPARGC1α*, reinforcing the conclusion that *PPARGC1α* is a critical mediator of sobetirome-induced protection against pulmonary fibrosis.

More importantly, sobetirome was well tolerated at the dose used in this study and did not induce weight loss, a common dose-limiting adverse effect associated with thyroid hormone therapy. However, caution regarding the long-term use of systemic thyromimetics remains warranted, as the related THRB-selective agonist eprotirome has been associated with cartilage toxicity following prolonged exposure [37]. To address this limitation and achieve more lung-selective drug delivery, we evaluated aerosolized administration of sobetirome and found that it retained robust therapeutic efficacy. Notably, aerosol-treated mice exhibited faster and more complete recovery of body weight, suggesting that systemic intraperitoneal administration may have exerted subtle metabolic effects that were not readily apparent. This interpretation is consistent with the known biology of thyroid hormone receptors: although cardiac effects are mediated primarily by THRA, the regulation of body weight and energy expenditure involves both THRA and THRB [55–57]. Consequently, limiting systemic exposure through inhaled delivery may preserve therapeutic efficacy while minimizing off-target metabolic effects [58]. Beyond minimizing systemic side effects, inhalational delivery offers several advantages for the treatment of lung diseases, including enhanced lung specificity, higher local drug concentrations, and reduced systemic exposure. This route also lowers the risk of off-target pharmacological interactions, an especially important consideration in patients with IPF, who are typically older and frequently receive multiple concomitant medications [59].

Our findings in human precision-cut lung slices provide important translational evidence that the antifibrotic effects of sobetirome extend beyond experimental animal models and are preserved in human lung tissue. Importantly, sobetirome was effective both in preventing the development of fibrosis in healthy human lung slices exposed to profibrotic stimuli and in reducing established fibrotic responses in lung tissue obtained from patients with IPF, supporting its potential therapeutic relevance in clinically advanced disease. Transcriptomic analyses further suggest that the effects of sobetirome are not limited to suppression of collagen synthesis but instead reflect broad remodeling of the fibrotic microenvironment. Notably, several genes consistently upregulated in IPF, including *COL1A1*, *COL3A1*, *POSTN*, *CTSK*, and *SFRP2* [60, 61], were markedly downregulated following sobetirome treatment, further supporting reversal of disease-associated transcriptional programs. In parallel, sobetirome simultaneously attenuated extracellular matrix production, myofibroblast activation, WNT and TGF-β signaling, cellular senescence, and cytoskeletal remodeling while promoting pathways associated with extracellular matrix turnover, basement membrane restoration, endothelial homeostasis, and immune regulation. These coordinated changes indicate that THRB activation promotes a multifaceted tissue repair program rather than targeting a single profibrotic pathway. Given the well-recognized limitations of animal models in predicting clinical efficacy, validation in human PCLS substantially strengthens the translational potential of sobetirome and supports further investigation of lung-targeted THRB agonism as a therapeutic strategy for IPF.

This study has several limitations. Although the bleomycin model is widely used, it does not fully recapitulate the chronic and progressive nature of human IPF [62]. While validation in human PCLS substantially strengthens the translational relevance of our findings, this *ex vivo* model cannot capture systemic drug effects or long-term tissue remodeling. In addition, although sobetirome was well tolerated in our study, its long-term safety and efficacy require evaluation in future preclinical and clinical studies. Nevertheless, the consistent antifibrotic effects observed across complementary *in vitro, in vivo*, and human *ex vivo* models provide strong support for the therapeutic potential of sobetirome as a selective THRB agonist in IPF.

## Materials and Methods

### Sex as a biological variable

In our study, we incorporated sex as a critical biological variable by including both male and female animals in *in vivo* assessments. All experimental groups were sex-balanced to ensure adequate statistical power to detect sex-dependent effects. Additionally, our biomarker discovery and mitochondrial DNA measurements will be performed after treatment on the fresh lung tissue slices from both sexes. We also tried to perform our *ex vivo* studies on the lung tissue slices from both sexes whenever possible

### Chemicals

Unless otherwise indicated, all materials were supplied by Sigma. Sobetirome was obtained from Emerson Resources, Inc. Antibodies and primers are summarized in supplementary **Table S3** and supplementary **Table S4**, respectively.

### Bleomycin model of lung fibrosis

The animal models of lung fibrosis were performed as previously described [18]. In brief, 9-12-week-old male C57BL/6 mice were purchased from Jackson Laboratories and anesthetized in an isoflurane inhalation chamber. Mice then either received 1.5 U/kg bleomycin (Hospira) in 50µl of 0,9% NaCl or 0,9% NaCl only. To assess therapeutic potential, sobetirome (300 µg/kg) was injected every other day, starting on day 8 until day 21. For inhalational administration, sobetirome was aerosolized (300 µg/kg, aerosol nebulizer (Omron)) every day from day 8 to 20. Mice were sacrificed on day 21. Mice with floxed *Ppargc1α* alleles and mice expressing *Col1a1-cre/ERT2* or *Sfptc-cre/ERT2* were purchased from Jackson Laboratories. To induce cell-specific KO, transgenic mice were treated with tamoxifen for 5 days before the start of experiments and every 72h thereafter as previously described [50].

### Hydroxyproline measurements

Hydroxyproline measurements were performed using the Biovision hydroxyproline kit as described previously. [18, 42]. In brief, after sacrificing the animals and perfusing the lung with PBS, the left lung lobe was separated and homogenized in ddH2O. An equal volume of 12 N HCl was added, and after incubation at 120°C for 3 hours, lysates were transferred to a 96-well plate, evaporated, and the measurements were performed using the Biovision hydroxyproline kit, according to the manufacturer’s instructions. Data are expressed as μg of hydroxyproline/left lung.

### Lung Function Study

Mice were subjected to lung function measurements by the forced oscillation technique on the FlexiVent® system (Scireq) as previously described [63, 64]. In brief, mice were anesthetized with urethane (1g/kg), and after the absence of pain responses, mice were connected to the ventilation unit and paralyzed with rocuronium bromide (1 mg/kg) [65]. Afterward, snapshot perturbations, forced oscillation perturbations, and ramp-volume regulated pressure-volume curves were performed at least three times per animal, and values were averaged[64]. Only measurements with at least two data points with a coefficient of determination above 0.95 were accepted.

### Histology

For all histological methods involving tissue, formaldehyde fixation, followed by paraffin-embedding, was performed, and 5µm sections were prepared on a microtome. Photomicrographs were evaluated based on a modified Ashcroft score in a blinded fashion on trichrome-stained sections [66]. All counting/grading was performed blinded on at least 5 randomly taken photomicrographs at 20x magnification. Positive cells for TUNEL staining and IF were obtained by using the ImageJ function [67].

### Immunohistochemistry/Immunofluorescence

In brief, sections were subjected to deparaffinization and heat-mediated antigen retrieval (citrate, pH=6, 2 x 5 minutes in the microwave), blocked with 2.5% secondary antibody host serum in PBS/0,3% Triton X, and stained overnight with primary antibody/antibodies as described previously [42].

Incubation with BLOXALL (Vector Laboratories) was performed for 10 minutes for IHC only. For immunofluorescence (IF), sections were then incubated with fluorochrome-labeled secondary antibodies raised in goat, while for immunohistochemistry, matching VECTASTAIN kits (Vector Laboratories) were used according to the manufacturer’s instructions. Immunofluorescence sections were subjected to TRUEVIEW autofluorescence quenching (Vector Laboratories) and mounted in a DAPI-containing mounting medium. For the same species’ immunofluorescence, a fluorescent alkaline phosphatase substrate (FastRed) was used first, and afterward, sections were microwaved, blocked, and subsequently stained with the second primary AB [42].

### In situ hybridization

In situ hybridization (ISH) was performed as previously described. [42]. In brief, sections were treated according to the manufacturer’s instructions (ACDbio) and incubated with probes against PPARGC1α and THRB, as well as a negative control. Afterward, detection was performed using the RNAScope Red kit (ACDbio), and subsequently, IF as described above was performed.

### Terminal nick end labeling

Terminal nick end labeling (TUNEL) was performed according to the manufacturer’s specifications using the in situ cell death detection kit. Tissue sections were rehydrated, washed, and incubated in permeabilization buffer (0,1% Triton X, 0,1% Citrate in ddH2O) for 8 minutes and washed afterward. Subsequently, slides were incubated with a reaction mixture containing fluorescently labeled dUDP and terminal deoxynucleotidyl transferase for 60 minutes at 37°C in a dark, humidified chamber. Afterward, sections were subjected to TRUEVIEW autofluorescence quenching (Vector Laboratories) and mounted in DAPI-containing mounting medium. If co-immunostaining was performed, a primary antibody was added during the 60-minute incubation, sections were washed, and incubated for 30 minutes with a fluorescently labeled secondary antibody. Cultured cells were fixed in 4% paraformaldehyde for 60 min at room temperature. After washing in PBS, they were permeabilized for 2 minutes on ice in permeabilization buffer (0,1% Triton X, 0,1% Citrate in ddH2O) and subsequently incubated with a reaction mixture containing fluorescently labeled dUDP and terminal deoxynucleotidyl transferase for 60 minutes at 37°C in a dark, humidified chamber. After the final washing step, cells were mounted in DAPI-containing mounting medium.

### Microscopy

For confocal images, an LSM710 confocal microscope (Zeiss) equipped with an argon and two-photon laser was used. Epifluorescence images were obtained on a Nikon Eclipse Ti microscope. Analysis was performed using ImageJ [67].

### Caspase 3/7 activity

The activity of caspase 3/7 was measured using the Caspase-Glo® 3/7 assay (Promega) according to the manufacturer’s instructions. The medium was removed from 96-well plates; 40µl of basal medium and 40 µl of reaction mix were added, and luminescence was measured on a microplate reader (Citation 3, BioTek). The obtained values were normalized to the respective vehicle.

### Cell culture

All incubators used were set to 5% CO2 and 37°C unless otherwise specified. Screening for mycoplasma contamination was performed regularly. Small airway epithelial cells were purchased from Lonza and cultured in a small airway basal medium supplemented with the SAGM kit [18]. All experiments were carried out in basal medium w/o supplements. For both apoptosis and measurements of mitochondrial function, cells were incubated with bleomycin (15 mU/ml) or vehicle (PBS) for 6h, supernatants were removed, and medium containing sobetirome (100 ng/ml) or vehicle (DMSO 0.01%) was added for 18 h, and respective measurements were performed. Reverse transfection was performed 48 hours before starting the experiments in medium w/o antibiotics using the Lipofectamine RNAiMAX kit (Thermo Fisher) according to the manufacturer’s instructions. siRNA Select and RNAiMAX Lipofectamine were diluted to a final concentration of 10 nM and 1:100, respectively, preincubated in OptiMEM for 20 minutes at RT, and 100µl of cell suspension, containing 90,000 cells/ml, was added. KD of selected genes was verified after 48 h. Fibroblasts derived from 4 different IPF patients were purchased from Lonza and cultured in FBM supplemented with an FGM-2 supplement kit. All experiments were carried out in a serum-free medium. For apoptosis, measurements of mitochondrial function, and electron microscopy, cells were treated with sobetirome (100 ng/ml), vehicle (DMSO 0.01%), or chloramphenicol (300 µM) for a total of 7 days[13] and were passaged the day before experiments were performed. 30,000 cells per well were seeded in 96-well plates/chamber slides of the same well size for both mitochondrial function and apoptosis measurements. For apoptosis, vehicle or FAS-ligand (300 ng/ml, R&D Systems) was added 24 h before measurements were performed. Normal human lung fibroblasts from Lonza were cultured in the same fashion and treated with recombinant human TGF (5 ng/ml, R&D Systems) before performing mitochondrial function tests. Reverse transfection was performed 48 hours before starting the experiments in medium w/o antibiotics and serum using the Lipofectamine RNAiMAX kit (Thermo Fisher) according to the manufacturer’s instructions. siRNA Select and RNAiMAX Lipofectamine were diluted to a final concentration of 10 nM and 1:100, respectively, preincubated in OptiMEM for 20 minutes at RT, and 100,000 cells were seeded. Because of the longer duration of experiments, cells were re-transfected on day 4 (final concentration of 1:300 for RNAiMAX and 40 nM siRNA Select) after initiation of treatment. Cells were harvested according to the experiments or verification of KD on day 7.

### Isolation of murine alveolar type 2 cells

Isolation of ATII cells was performed as previously described with some modifications [18, 68, 69]. In brief, mice were anesthetized, the aorta was cut, and the lungs were perfused with PBS. Afterward, 1 ml of dispase II (50 mg/ml in DMEM supplemented with antibiotics/antimycotics) was injected into the lungs via a cannula, followed by 0.5 ml of 1% low-melt agarose. After 45 min of incubation at room temperature, lungs were minced in DNase I (0,1 mg/ml in DMEM supplemented with antibiotics/antimycotics), and a single cell suspension was prepared by filtering and centrifuging. Red blood cells were lysed using ACK lysis buffer, and after additional centrifugation, the supernatant was discarded, and cells were incubated with FITC-labelled antibodies against CD11b, F4/80, CD11c, CD16/32, CD45, and CD19 for 10 minutes at 4 °C in the dark. Cells were washed and gated for single cells and FITC- and side-scatter high cells (see Supplementary Figure S3). Afterwards, cells were counted, and 60,000 (mitochondrial function) or 10,000 (apoptosis) cells per well were seeded. For both apoptosis and measurements of mitochondrial function, cells were incubated with bleomycin (15 mU/ml) or vehicle (PBS) for 6 h, supernatants were removed, and medium containing sobetirome (100 ng/ml) or vehicle (DMSO 0.01%) was added for 18 h, and respective measurements were performed.

### Mitochondrial function test

Oxygen consumption rate (OCR) and extracellular acidification rate (ECAR) were measured using the Seahorse XFe96 analyzer (Agilent) as previously described for both epithelial cells [18] and fibroblasts[13]. In brief, cells were incubated for 60 minutes in XF DMEM (pH=7.4, with 5mM HEPES) containing 2mM glutamine, 10mM glucose, and 1 mM pyruvate (all Agilent) in an incubator set to 37°C and 0% CO_2_. Immediately afterward, measurements were started, and oligomycin (1.5 µM final concentration), FCCP (1.0 µM final concentration), and rotenone/antimycin A (0.5 µM final concentration) were added as indicated in the graphs.

### Western blot

Western blotting was performed as previously described [18, 68]. In brief, cells or tissue were lysed in lysis buffer (M-PER or T-PER, both Thermo Fisher) supplemented with protease and phosphatase inhibitor cocktail (Abcam), and protein concentration was determined using the BCA kit (Thermo Fisher). 10 µg of protein was loaded per lane (4-20% gel, Bio-Rad) and transferred to a PVDF membrane (Trans-Blot Turbo, Bio-Rad). After blocking (5% BSA/dry milk, AmericanBio), membranes were incubated overnight with primary antibody at 4°C, washed, and subsequently, HRP-conjugated secondary antibody was added. Signals were detected using Clarity ECL substrate (Bio-Rad) and a Chemidoc MP Imaging system. Densitometric analysis was performed using ImageLab.

### Collagen ELISA

Enzyme-linked immunosorbent assay (ELISA) for measuring Collagen 1 alpha 1 (COL1A1)/procollagen 1 alpha 1 was performed using the Duo set kit from Bio-Techne according to the manufacturer’s instructions. Values were obtained by subtracting absorption at 540 nm from that at 450 nm on a microplate reader. Data were normalized to protein concentrations in the supernatants.

### Mitobiogenesis

Mitobiogenesis was measured using a commercially available in-cell ELISA directed against the subunit I of complex IV (COXI, mitochondrial DNA encoded) and the 70 kDa subunit of Complex II (SDH-A, nuclear DNA encoded) according to the manufacturer’s instructions (Abcam). Mitobiogenesis is expressed as the ratio of COXI/SDH-A, and chloramphenicol, a known inhibitor of mitochondrial protein synthesis and mitobiogenesis, was used as a negative control [47, 70].

### RNA extraction and qPCR

Total RNA was extracted from 30 – 50 mg of frozen lung tissue in 700 μL of Qiazol (Lysis buffer, Qiagen) using the miRNeasy Mini Kit (Qiagen) according to the manufacturer’s instructions. The purity of the RNA was verified using NanoDrop at 260 nm. Gene expression was determined by TaqMan® (Life Technologies) according to the manufacturer’s instructions. Glyceraldehyde-3-Phosphate Dehydrogenase (Gapdh) was employed as an internal standard control, and the specific primers and probes were all obtained from Life Technologies (Thermo Scientific Inc.). Primers used were Relative gene expressions normalized to a value of 1.0 for the unstimulated control group or vehicle-treated cells. Control reactions without RNA were done as a negative control. Fold change was calculated by taking the means of the controls as the baseline.

### Transmission electron microscopy

Electron microscopy was performed as previously described [71]. Cells were grown in the presence or absence of sobetirome for 7 days and fixed using Karnovsky’s fixation protocol. After processing for TEM, images were acquired using a JEM 1011 TEM microscope.

### Human Precision Cut Lung Slices (hPCLS)

Fresh human lungs, derived from patients with IPF or healthy donors, were cultured and used in two independent experiments [36, 72, 73]. The human PCLSs (hPCLS) harvested from healthy donors were cultured and treated with a medium containing a profibrotic cocktail (FC) (containing TGF-β, PDGF-AB [platelet-derived growth factor-AB], lysophosphatidic acid, TNF-α [tumor necrosis factor-α]) or a control cocktail [36]. The hPCLSs isolated from patients with IPF, as well as individual PCLSs treated with FC or a control cocktail supplemented with sobetirome (100 ng/m) or vehicle for 5 days, were used for advanced analysis. In brief, low-melting-agarose-instilled lobes were separated into tissue blocks, cut using a vibratome (300µm thick slices), and cultured in DMEM/F12, supplemented with antibiotics/antimycotics in 24-well plates. After 24 hours, slices were washed, and sobetirome or vehicle was added. After 120 h, the tissue was used for RNA isolation or embedding.

### RNA-Sequencing. RNAseq. Library Preparation, Sequencing, and Data Processing

Total RNA was extracted with the miRNeasy Mini Kit (Qiagen, 217004). RNA quantity and purity were assessed using a NanoDrop at 260nm as well as an Agilent 2100 Bioanalyzer. cDNA libraries were prepared with the KAPA Stranded mRNA-seq Kit and subsequently assessed with an Agilent TapeStation analyzer. Libraries were sequenced on a NovaSeq 6000 (Illumina) in a 2×100 paired-end format, aiming for a minimum of 25 million reads per sample. Reads were mapped to either the human (GRCh38) or mouse (GRCm39) genome reference with the alignment tool STAR (version 2.6.1d). ***RNA-seq Analysis:*** A similar analysis strategy was applied to both mouse and IPF PCLS RNASeq datasets. Analysis was performed in R (version 4.3.1) using the package DESeq2 (version 1.42.1). Genes with fewer than 100 total fragments across samples were discarded. Hypothesis testing was performed with DeSeq2’s default Wald tests under multifactor design conditions. The overlap of significant results from each test was visualized with the R package eulerr (version 7.0.2). The heatmap data was normalized with DESeq2’s variance-stabilizing transformation, then values across each gene were min-max normalized to fit the range 0 to 1. Genes in each heatmap region were clustered and organized using the R package seriation (version 1.5.7), where a distance matrix of the normalized values was re-ordered by optimal leaf ordering. The resulting dendrograms were clustered with the R package dendextend’s (version 1.17.1) *cutree* implementation and a *k* parameter of 2. Genes belonging to each cluster were then subject to gene set enrichment testing for Gene Ontology Biological Process terms (version 2025-03-16) for either Homo sapiens or Mus muscalis, using the PANTHER over-representation test (Fisher test option; version 20240807). Differentially expressed genes (DEGs) identified from bulk RNA sequencing were analyzed using Ingenuity Pathway Analysis (IPA; QIAGEN) to identify enriched canonical pathways and upstream regulatory networks. Gene lists, including corresponding log₂ fold change and statistical significance values, were uploaded into IPA and filtered using a significance threshold of *P* < 0.05. Canonical pathway enrichment was determined using a right-tailed Fisher’s exact test, which assesses the likelihood that the association between the input gene set and a given pathway occurs by chance. Pathways meeting the significance cutoff were considered enriched. Overlap between canonical pathways was further evaluated based on the proportion of shared genes relative to the total number of genes within each pathway, allowing identification of interconnected signaling networks. Upstream regulator analysis was performed within IPA to predict activation or inhibition of key transcriptional regulators based on the directionality of gene expression changes, providing insight into signaling nodes modulated by sobetirome.

### Statistics

Statistical analysis was performed using GraphPad Prism 9 and R for sequencing data. Where applicable, data were analyzed for normality using the D’Agostino-Pearson test. Two groups were compared using Student’s T-test or Mann-Whitney U test if the data did not follow a normal distribution. For three or more groups, one-way ANOVA, followed by Holm-Sidak’s multiple-comparison test, was used unless indicated otherwise, and the Kruskal-Wallis test, followed by Dunn’s test, was performed for non-parametric data. For time-course experiments (i.e., weight, OCR, pressure-volume curves), two-way ANOVA for repeated measurements followed by Holm-Sidak’s multiple-comparison test was used.

### Study approval

*Animal studies:* All animal studies were conducted in accordance with the NIH guidelines, and approval was obtained from the institutional committee at Yale University (2023-11592). *Human Studies:* Age- and sex-matched healthy lungs were from rejected donor lung organs provided by the National Disease Research Interchange (NDRI). The IPF lung explants were provided by Baylor College of Medicine. All human lung samples were provided without any identifying information, and the study was determined as non-human subject research by the Yale IRB committee (protocol no. 2000024530) and Baylor College of Medicine (IRB H-46832).

## Data availability

All data, code, and materials used in the analysis are available to any researcher for purposes of reproducing or extending the analysis. All raw count expression data from the *in vivo* and *ex vivo* studies were deposited in the Gene Expression Omnibus (GEO), and the accession number is GSE335112.

## Author contributions

F.A. and N.K. conceptualized, designed, and supervised the study. All *in vitro* studies were performed by T.B, S.D., and K.R. All *in vivo* studies were performed by T.B, S.D., and F.A. The human precision-cut lung slices for *ex vivo* studies were provided by F.F. and I.R., and all experiments and analysis on human PCLS were performed by T.B, F.A., R.R., J.K., M.S., L.L., K.V., A.J., and S.A. Transcriptomic data were processed, curated, and visualized by T.A., A.J., C.C., and F.N. G.D. coordinated and prepared the necessary materials for the experiments. The manuscript was drafted and edited by T.B., N.K., and F.A., and was reviewed and edited by all other authors.

## Acknowledgments

We want to express our gratitude to Rebecca Cardone and Xiaojian Zhao at the Chemical Metabolism Core at Yale School of Medicine for their expert assistance with Seahorse experiments. Similarly, we thank the Electron Microscopy facility at the Center for Cellular and Molecular Imaging (CCMI) at Yale School of Medicine for all the help they provided for tissue processing and imaging, as well as the Yale Center for Genome Analysis (YGCA) for their help in performing RNASeq.

## References

1. Raghu, G., et al., Idiopathic Pulmonary Fibrosis (an Update) and Progressive Pulmonary Fibrosis in Adults: An Official ATS/ERS/JRS/ALAT Clinical Practice Guideline. Am J Respir Crit Care Med, 2022. 205(9): p. e18–e47.

2. Raghu, G. and T.R. Fleming, Moving forward in IPF: lessons learned from clinical trials. Lancet Respir Med, 2024. 12(8): p. 583–585.

3. Gomer, R.H., New approaches to modulating idiopathic pulmonary fibrosis. Curr Allergy Asthma Rep, 2013. 13(6): p. 607–12.

4. King, T.E., Jr., et al., A phase 3 trial of pirfenidone in patients with idiopathic pulmonary fibrosis. N Engl J Med, 2014. 370(22): p. 2083–92.

5. George, G., U. Vaid, and R. Summer, Therapeutic advances in idiopathic pulmonary fibrosis. Clin Pharmacol Ther, 2016. 99(1): p. 30–2.

6. Richeldi, L., et al., Nerandomilast in Patients with Idiopathic Pulmonary Fibrosis. N Engl J Med, 2025. 392(22): p. 2193–2202.

7. Barbas-Filho, J.V., et al., Evidence of type II pneumocyte apoptosis in the pathogenesis of idiopathic pulmonary fibrosis (IFP)/usual interstitial pneumonia (UIP). J Clin Pathol, 2001. 54(2): p. 132–8.

8. Thannickal, V.J. and J.C. Horowitz, Evolving concepts of apoptosis in idiopathic pulmonary fibrosis. Proc Am Thorac Soc, 2006. 3(4): p. 350–6.

9. Kis, K., X. Liu, and J.S. Hagood, Myofibroblast differentiation and survival in fibrotic disease. Expert Rev Mol Med, 2011. 13: p. e27.

10. Redente, E.F., et al., Loss of Fas signaling in fibroblasts impairs homeostatic fibrosis resolution and promotes persistent pulmonary fibrosis. JCI Insight, 2020. 6(1).

11. Hagimoto, N., et al., Induction of apoptosis and pulmonary fibrosis in mice in response to ligation of Fas antigen. Am J Respir Cell Mol Biol, 1997. 17(3): p. 272–8.

12. Warsinske, H.C., et al., Computational Modeling Predicts Simultaneous Targeting of Fibroblasts and Epithelial Cells Is Necessary for Treatment of Pulmonary Fibrosis. Front Pharmacol, 2016. 7: p. 183.

13. Ryu, C., et al., Extracellular Mitochondrial DNA Is Generated by Fibroblasts and Predicts Death in Idiopathic Pulmonary Fibrosis. Am J Respir Crit Care Med, 2017. 196(12): p. 1571–1581.

14. Rangarajan, S., et al., Metformin reverses established lung fibrosis in a bleomycin model. Nat Med, 2018. 24(8): p. 1121–1127.

15. Parimon, T., et al., Alveolar Epithelial Type II Cells as Drivers of Lung Fibrosis in Idiopathic Pulmonary Fibrosis. Int J Mol Sci, 2020. 21(7).

16. Bueno, M., et al., PINK1 deficiency impairs mitochondrial homeostasis and promotes lung fibrosis. J Clin Invest, 2015. 125(2): p. 521–38.

17. Patel, A.S., et al., Epithelial cell mitochondrial dysfunction and PINK1 are induced by transforming growth factor-beta1 in pulmonary fibrosis. PLoS One, 2015. 10(3): p. e0121246.

18. Yu, G., et al., Thyroid hormone inhibits lung fibrosis in mice by improving epithelial mitochondrial function. Nat Med, 2018. 24(1): p. 39–49.

19. Bianco, A.C. and R.R. da Conceicao, The Deiodinase Trio and Thyroid Hormone Signaling. Methods Mol Biol, 2018. 1801: p. 67–83.

20. Singh, B.K. and P.M. Yen, A clinician’s guide to understanding resistance to thyroid hormone due to receptor mutations in the TRalpha and TRbeta isoforms. Clin Diabetes Endocrinol, 2017. 3: p. 8.

21. Harper, M.E. and E.L. Seifert, Thyroid hormone effects on mitochondrial energetics. Thyroid, 2008. 18(2): p. 145–56.

22. Weitzel, J.M., K.A. Iwen, and H.J. Seitz, Regulation of mitochondrial biogenesis by thyroid hormone. Exp Physiol, 2003. 88(1): p. 121–8.

23. Lebon, V., et al., Effect of triiodothyronine on mitochondrial energy coupling in human skeletal muscle. J Clin Invest, 2001. 108(5): p. 733–7.

24. Fernandez-Marcos, P.J. and J. Auwerx, Regulation of PGC-1alpha, a nodal regulator of mitochondrial biogenesis. Am J Clin Nutr, 2011. 93(4): p. 884S–90.

25. Eghtedari, B. and R. Correa, Levothyroxine, in StatPearls. 2021: Treasure Island (FL).

26. Scanlan, T.S., Sobetirome: a case history of bench-to-clinic drug discovery and development. Heart Fail Rev, 2010. 15(2): p. 177–82.

27. Pan, X., et al., TRbeta activation confers AT2-to-AT1 cell differentiation and anti-fibrosis during lung repair via KLF2 and CEBPA. Nat Commun, 2024. 15(1): p. 8672.

28. Ajayi, I.O., et al., X-linked inhibitor of apoptosis regulates lung fibroblast resistance to Fas-mediated apoptosis. Am J Respir Cell Mol Biol, 2013. 49(1): p. 86–95.

29. Cooley, J.C., et al., Inhibition of antiapoptotic BCL-2 proteins with ABT-263 induces fibroblast apoptosis, reversing persistent pulmonary fibrosis. JCI Insight, 2023. 8(3).

30. Irrcher, I., et al., Thyroid hormone (T3) rapidly activates p38 and AMPK in skeletal muscle in vivo. J Appl Physiol (1985), 2008. 104(1): p. 178–85.

31. Sinha, R.A., et al., Thyroid hormone induction of mitochondrial activity is coupled to mitophagy via ROS-AMPK-ULK1 signaling. Autophagy, 2015. 11(8): p. 1341–57.

32. Adams, T.S., et al., Single-cell RNA-seq reveals ectopic and aberrant lung-resident cell populations in idiopathic pulmonary fibrosis. Sci Adv, 2020. 6(28): p. eaba1983.

33. Richter, U., et al., A mitochondrial ribosomal and RNA decay pathway blocks cell proliferation. Curr Biol, 2013. 23(6): p. 535–41.

34. McKee, E.E., et al., Inhibition of mammalian mitochondrial protein synthesis by oxazolidinones. Antimicrob Agents Chemother, 2006. 50(6): p. 2042–9.

35. Ahangari, F., et al., Saracatinib Is a Potential Novel Therapeutic for Pulmonary Fibrosis. American Journal of Respiratory and Critical Care Medicine, 2020. 201.

36. Alsafadi, H.N., et al., An ex vivo model to induce early fibrosis-like changes in human precision-cut lung slices. Am J Physiol Lung Cell Mol Physiol, 2017. 312(6): p. L896–L902.

37. Lammel Lindemann, J. and P. Webb, Sobetirome: the past, present and questions about the future. Expert Opin Ther Targets, 2016. 20(2): p. 145–9.

38. Wang, L., et al., Single-Cell RNA-seq Provides New Insights into Therapeutic Roles of Thyroid Hormone in the Idiopathic Pulmonary Fibrosis. Am J Respir Cell Mol Biol, 2023.

39. Lama, V., et al., Prostaglandin E2 synthesis and suppression of fibroblast proliferation by alveolar epithelial cells is cyclooxygenase-2-dependent. Am J Respir Cell Mol Biol, 2002. 27(6): p. 752–8.

40. Desai, T.J., D.G. Brownfield, and M.A. Krasnow, Alveolar progenitor and stem cells in lung development, renewal and cancer. Nature, 2014. 507(7491): p. 190–4.

41. Lehmann, M., et al., Differential effects of Nintedanib and Pirfenidone on lung alveolar epithelial cell function in ex vivo murine and human lung tissue cultures of pulmonary fibrosis. Respir Res, 2018. 19(1): p. 175.

42. Barnthaler, T., et al., Inhibiting eicosanoid degradation exerts antifibrotic effects in a pulmonary fibrosis mouse model and human tissue. J Allergy Clin Immunol, 2019.

43. Wan, Z., et al., Evidence for the role of AMPK in regulating PGC-1 alpha expression and mitochondrial proteins in mouse epididymal adipose tissue. Obesity (Silver Spring), 2014. 22(3): p. 730–8.

44. Marquez-Jurado, S., et al., Mitochondrial levels determine variability in cell death by modulating apoptotic gene expression. Nat Commun, 2018. 9(1): p. 389.

45. Tang, D., et al., Metformin facilitates BG45induced apoptosis via an antiWarburg effect in cholangiocarcinoma cells. Oncol Rep, 2018. 39(4): p. 1957–1965.

46. Xie, N., et al., Glycolytic Reprogramming in Myofibroblast Differentiation and Lung Fibrosis. Am J Respir Crit Care Med, 2015. 192(12): p. 1462–74.

47. Kao, L.P., D. Ovchinnikov, and E. Wolvetang, The effect of ethidium bromide and chloramphenicol on mitochondrial biogenesis in primary human fibroblasts. Toxicol Appl Pharmacol, 2012. 261(1): p. 42–9.

48. Caporarello, N., et al., PGC1alpha repression in IPF fibroblasts drives a pathologic metabolic, secretory and fibrogenic state. Thorax, 2019. 74(8): p. 749–760.

49. Marrif, H., et al., Temperature homeostasis in transgenic mice lacking thyroid hormone receptor-alpha gene products. Endocrinology, 2005. 146(7): p. 2872–84.

50. Tsoyi, K., et al., CD148 Deficiency in Fibroblasts Promotes the Development of Pulmonary Fibrosis. Am J Respir Crit Care Med, 2021. 204(3): p. 312–325.

51. Tsukui, T., et al., Collagen-producing lung cell atlas identifies multiple subsets with distinct localization and relevance to fibrosis. Nat Commun, 2020. 11(1): p. 1920.

52. Hung, C., et al., Role of lung pericytes and resident fibroblasts in the pathogenesis of pulmonary fibrosis. Am J Respir Crit Care Med, 2013. 188(7): p. 820–30.

53. Lu, S., et al., Smooth muscle-derived progenitor cell myofibroblast differentiation through KLF4 downregulation promotes arterial remodeling and fibrosis. JCI Insight, 2020. 5(23).

54. Jenkins, R.G., et al., An Official American Thoracic Society Workshop Report: Use of Animal Models for the Preclinical Assessment of Potential Therapies for Pulmonary Fibrosis. Am J Respir Cell Mol Biol, 2017. 56(5): p. 667–679.

55. Grover, G.J., et al., Selective thyroid hormone receptor-beta activation: a strategy for reduction of weight, cholesterol, and lipoprotein (a) with reduced cardiovascular liability. Proc Natl Acad Sci U S A, 2003. 100(17): p. 10067–72.

56. Wikstrom, L., et al., Abnormal heart rate and body temperature in mice lacking thyroid hormone receptor alpha 1. EMBO J, 1998. 17(2): p. 455–61.

57. Bochukova, E., et al., A mutation in the thyroid hormone receptor alpha gene. N Engl J Med, 2012. 366(3): p. 243–9.

58. Borghardt, J.M., C. Kloft, and A. Sharma, Inhaled Therapy in Respiratory Disease: The Complex Interplay of Pulmonary Kinetic Processes. Can Respir J, 2018. 2018: p. 2732017.

59. Khor, Y.H., I. Glaspole, and N.S.L. Goh, Therapeutic burden in interstitial lung disease: Lessons to learn. Respirology, 2019. 24(6): p. 566–571.

60. McDonough, J.E., et al., Transcriptional regulatory model of fibrosis progression in the human lung. JCI Insight, 2019. 4(22).

61. Buhling, F., et al., Pivotal role of cathepsin K in lung fibrosis. Am J Pathol, 2004. 164(6): p. 2203–16.

62. Brazee, P., et al., Peeling Back the Layers of the Bleomycin Model of Lung Fibrosis: Lessons Learned, Factors to Consider, and Future Directions. Semin Respir Crit Care Med, 2025. 46(4): p. 330–346.

63. Frei, R.B., et al., Cannabinoid receptor 2 augments eosinophil responsiveness and aggravates allergen-induced pulmonary inflammation in mice. Allergy, 2016. 71(7): p. 944–56.

64. Vanoirbeek, J.A., et al., Noninvasive and invasive pulmonary function in mouse models of obstructive and restrictive respiratory diseases. Am J Respir Cell Mol Biol, 2010. 42(1): p. 96–104.

65. Lovgren, A.K., et al., COX-2-derived prostacyclin protects against bleomycin-induced pulmonary fibrosis. Am J Physiol Lung Cell Mol Physiol, 2006. 291(2): p. L144–56.

66. Hubner, R.H., et al., Standardized quantification of pulmonary fibrosis in histological samples. Biotechniques, 2008. 44(4): p. 507–11, 514-7.

67. Schneider, C.A., W.S. Rasband, and K.W. Eliceiri, NIH Image to ImageJ: 25 years of image analysis. Nat Methods, 2012. 9(7): p. 671–5.

68. Barnthaler, T., et al., The Role of PGE2 in Alveolar Epithelial and Lung Microvascular Endothelial Crosstalk. Sci Rep, 2017. 7(1): p. 7923.

69. Gereke, M., et al., Flow cytometric isolation of primary murine type II alveolar epithelial cells for functional and molecular studies. J Vis Exp, 2012(70).

70. Lipton, J.H. and W.C. McMurray, Mitochondrial biogenesis in cultured animal cells. I. Effect of chloramphenicol on morphology and mitochondrial respiratory enzymes. Biochim Biophys Acta, 1977. 477(3): p. 264–72.

71. Ahangari, F., et al., microRNA-33 deficiency in macrophages enhances autophagy, improves mitochondrial homeostasis, and protects against lung fibrosis. JCI Insight, 2023. 8(4).

72. Ahangari, F., et al., Saracatinib, a Selective Src Kinase Inhibitor, Blocks Fibrotic Responses in Preclinical Models of Pulmonary Fibrosis. Am J Respir Crit Care Med, 2022. 206(12): p. 1463–1479.

73. Liu, G., et al., Use of precision cut lung slices as a translational model for the study of lung biology. Respir Res, 2019. 20(1): p. 162.

