## Supplementary material for "Sobetirome, a thyroid hormone receptor beta agonist, is a potential therapeutic agent for pulmonary fibrosis": Suplemental figures and tables

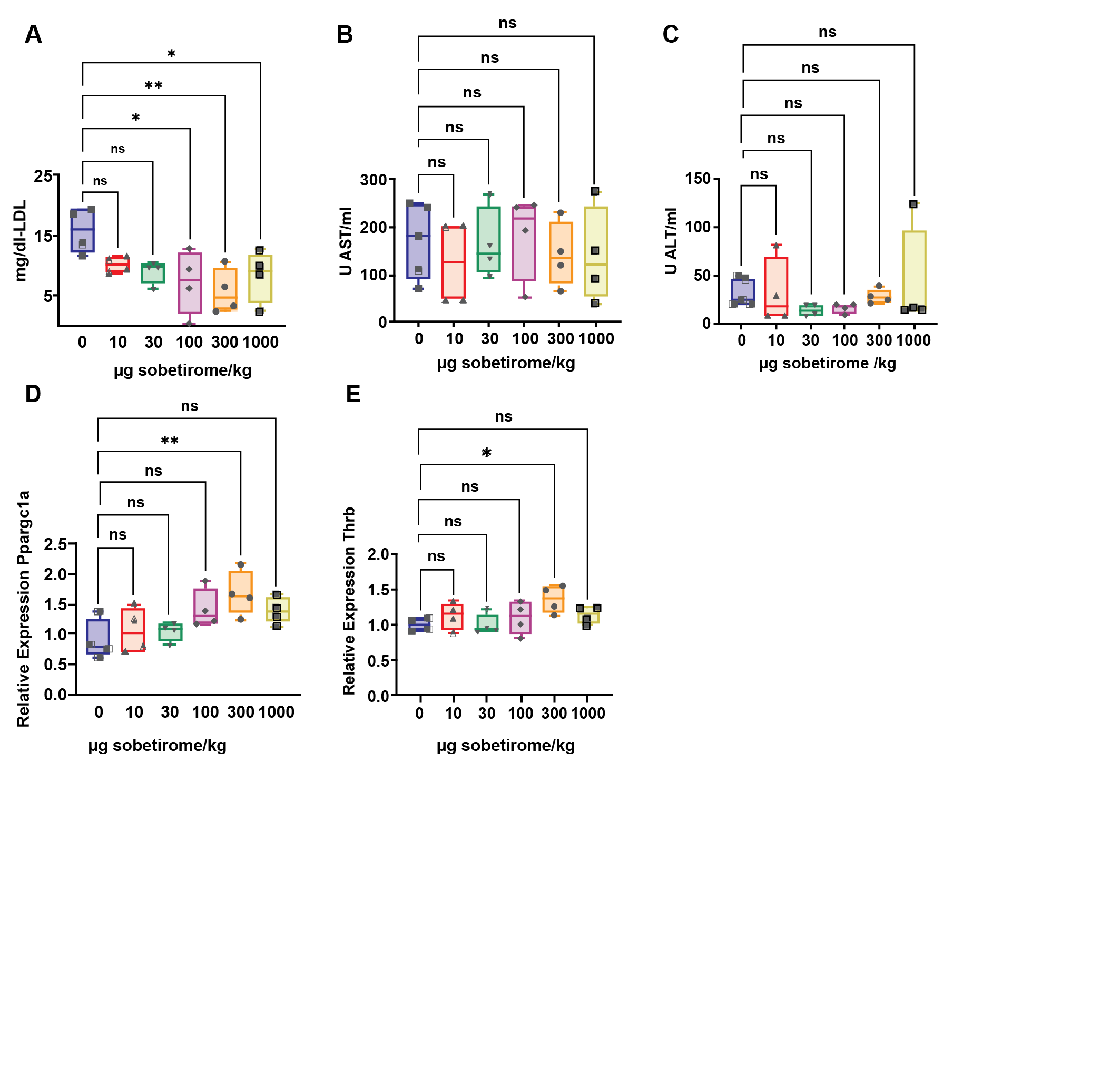
**Supplementary Figure S1.**

**Supplementary Figure S1. Intraperitoneal administration of sobetirome decreases LDL and increases *PPARGC1a* and *THRB* but does not affect ALT or AST.** Mice were treated every other day for 14 days at the indicated dose and were subsequently sacrificed. **(A)** LDL, **(B)** AST, and **(C)** ALT were measured in plasma via ELISA (LDL) and activity assays (ALT and AST), respectively. **(D)** *PPARGC1a* and **(E)** *THRB* relative expressions were determined by qPCR in the lungs. (n=4-5), ns= non-significant, *=p<0.05, **=p<0.01.

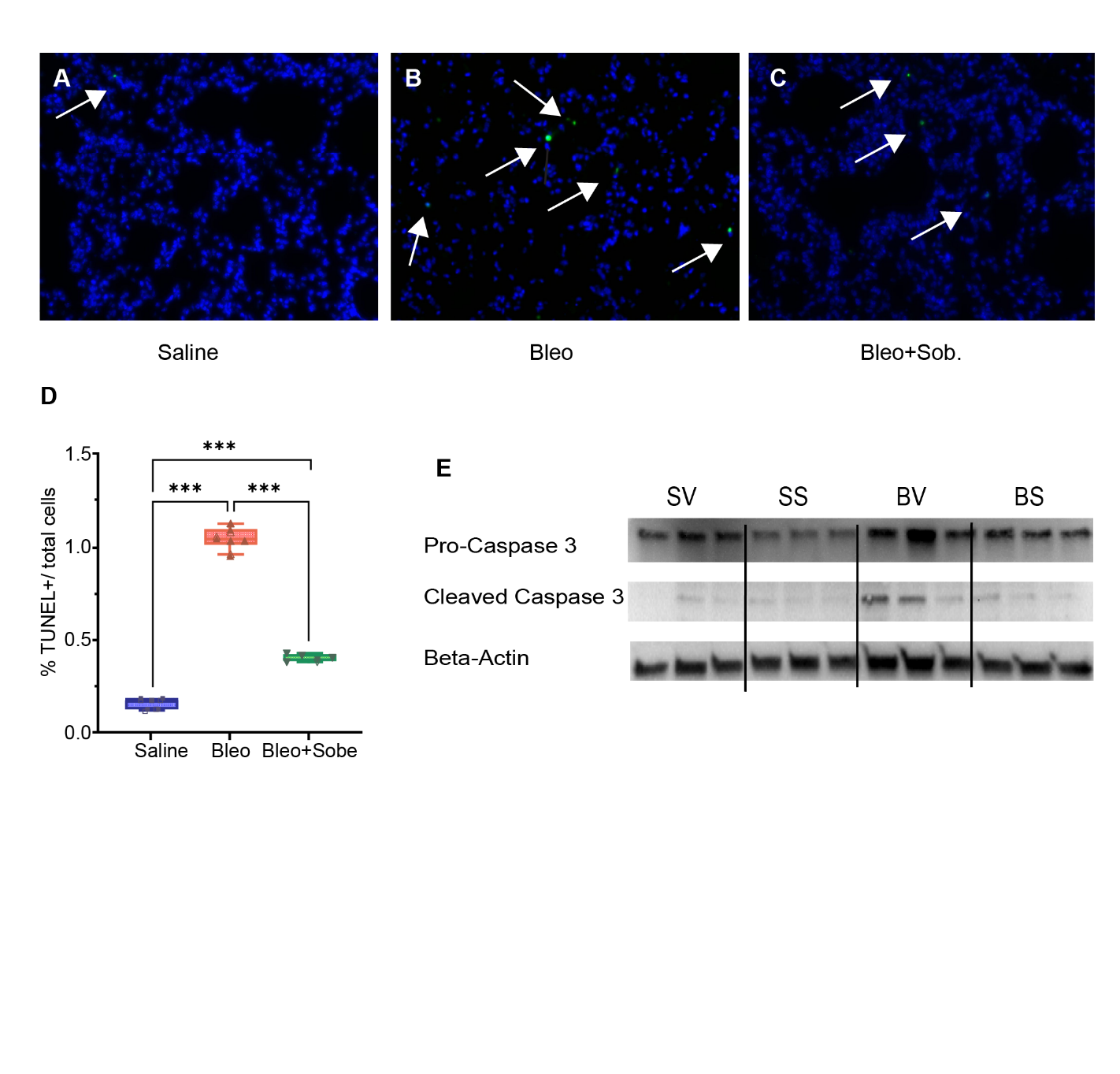
**Supplementary Figure S2.**

**Supplementary Figure S2. Sobetirome decreases bleomycin-induced apoptosis.** Lungs from **(A)** vehicle-, **(B)** bleomycin-, and **(C)** bleomycin + sobetirome-treated mice were stained with TUNEL reagent and counterstained with DAPI. **(D)** shows quantification of positive-staining cells. **(E)** shows a representative Western blot of the respective groups and indicated targets. The samples shown are the same ones as in Fig. 1L (beta-actin is the same). One-way ANOVA was performed for D. ***=p<0.001; arrows show positive cells.

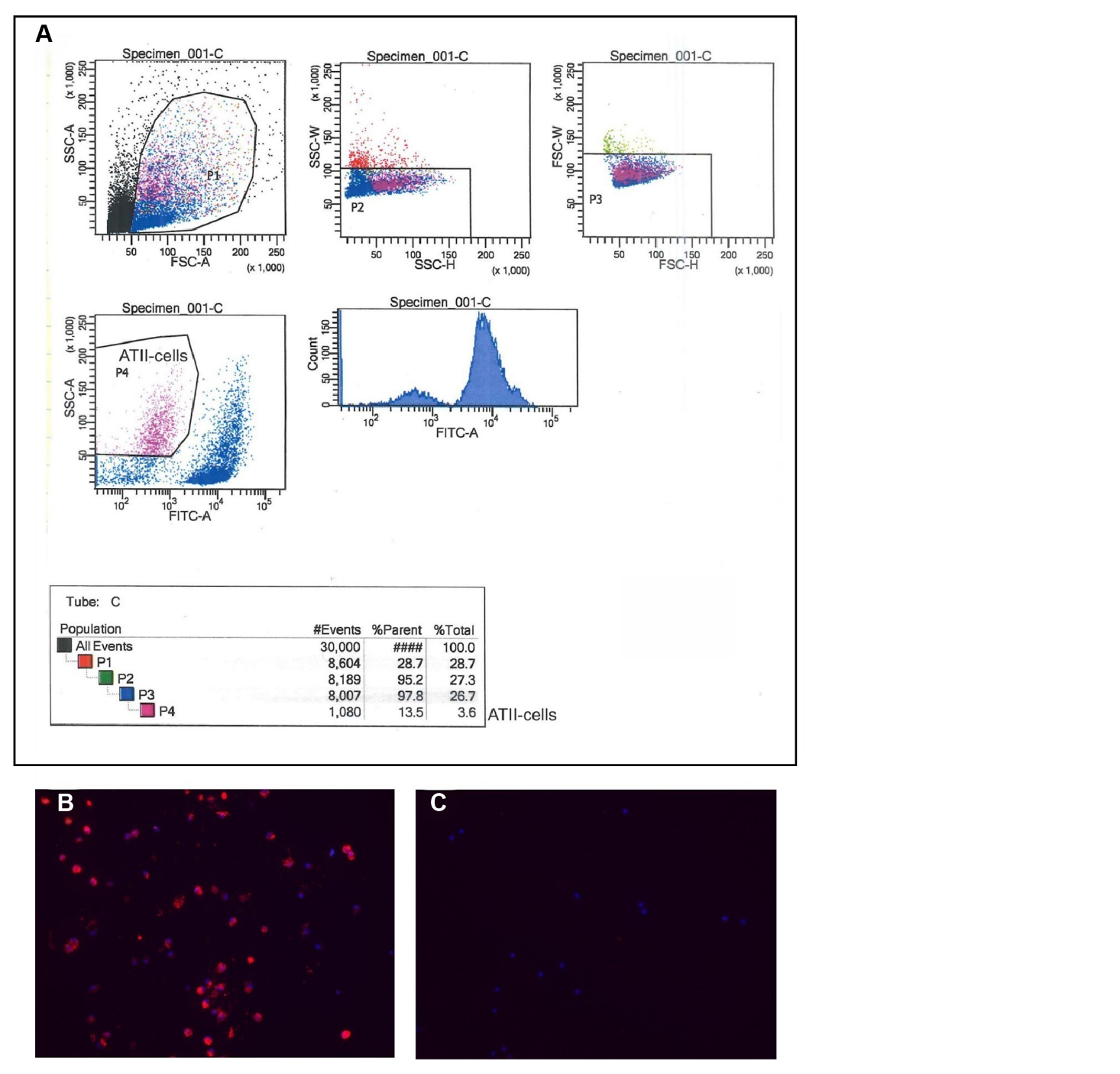
**Supplementary Figure S3.**

**Supplementary Figure S3. Alveolar type II cells were isolated from murine lungs. (A)** shows the gating strategy, with side scatter high; FITC (CD16/32, CD45, CD11b, CD11c, F4/80, CD19)-negative cells are the desired cell population. **(B)** After 24 h in culture, cells were stained with an antibody directed against pro-SPC (red) or (C) without primary antibody.

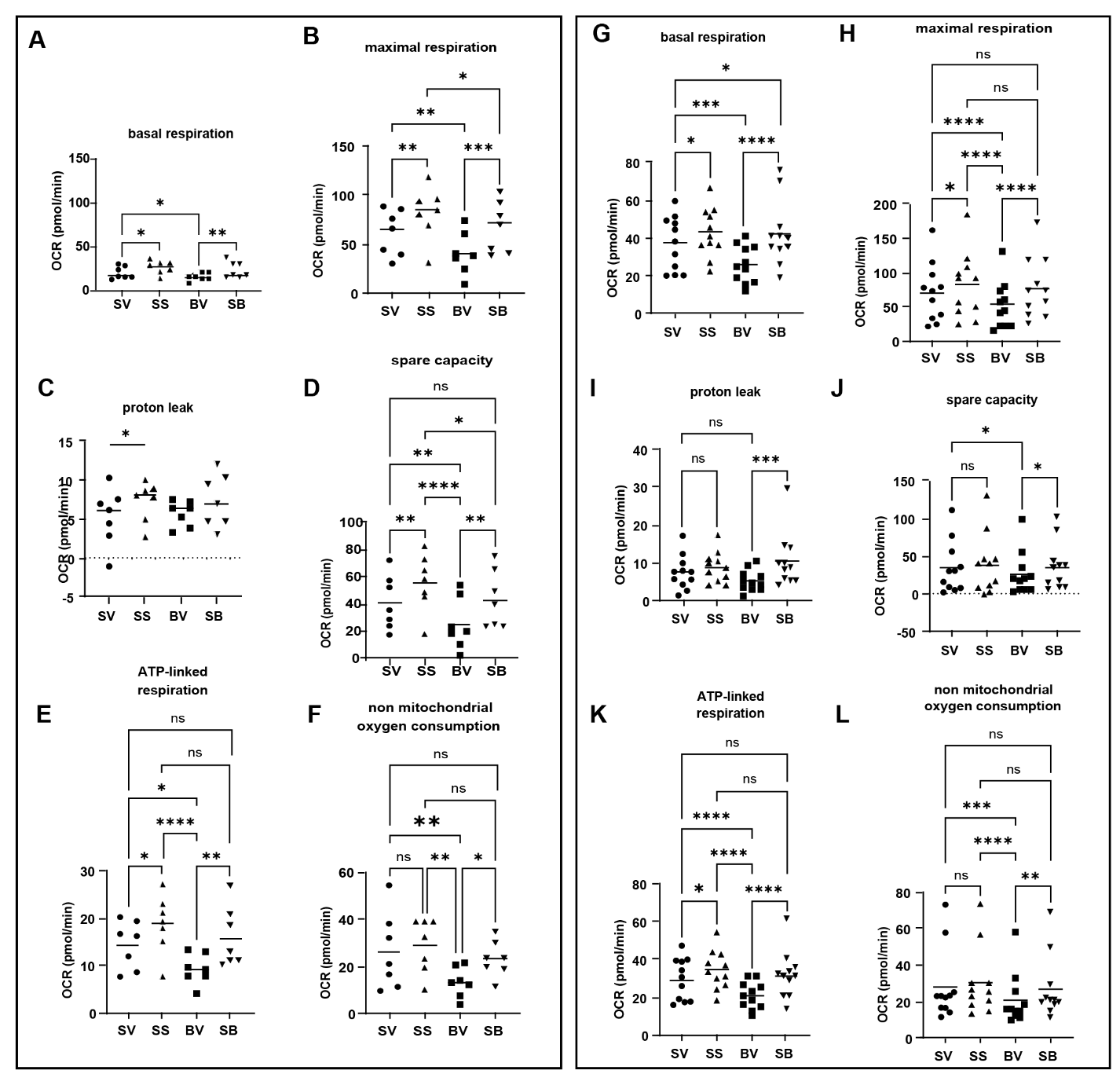
**Supplementary Figure S4.**

**Supplementary Figure S4. Sobetirome improves a variety of mitochondrial readouts in lung epithelial cells.** (**A-F**) Isolated murine ATII cells and (**G-L**) small airway epithelial cells were stimulated with bleomycin (B) or saline (S), supernatant was removed, and cells were subsequently treated with vehicle (V) or sobetirome (S). Oxygen consumption rate was measured using the Seahorse device, and the indicated parameters were calculated. One-way ANOVA was performed. *=p<0.05, **=p<0.01, ***=p<0.001, ****= p<0.0001.

**Supplementary Figure S5**
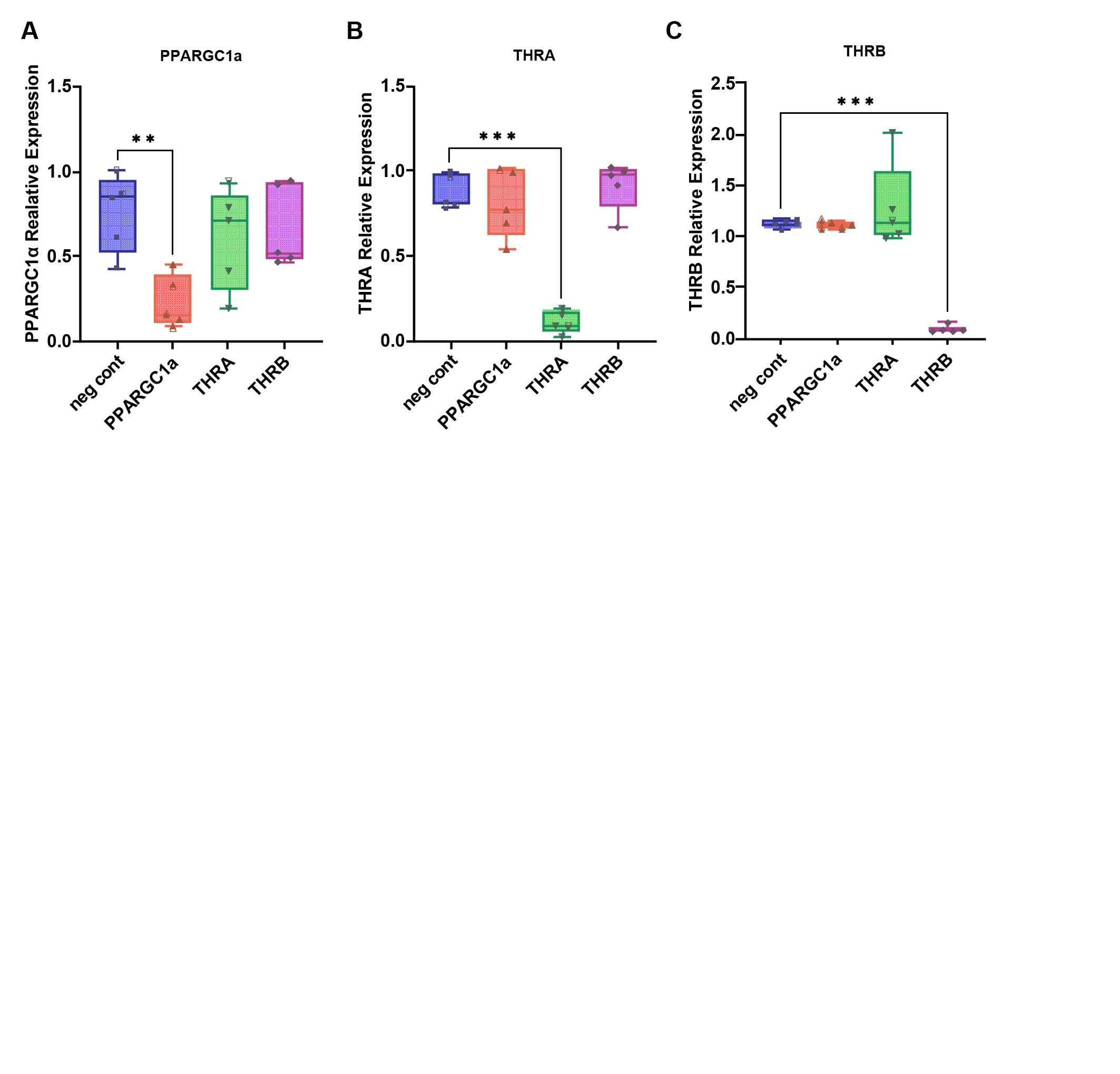
**.**

**Supplementary Figure S5. SAECs were transfected with negative control siRNA or siRNA targeted to *PPARGC1α*, *THRA,* or *THRB.*** Levels of **(A)** PPARGC1α, **(B)** *THRA,* and **(C)** *THRB* were assessed after 48 h and are provided as normalized to a non-transfected control. Statistical analysis was done by One-Way ANOVA, **=p<0.01, ***=p<0.001, ***=p<0.001.

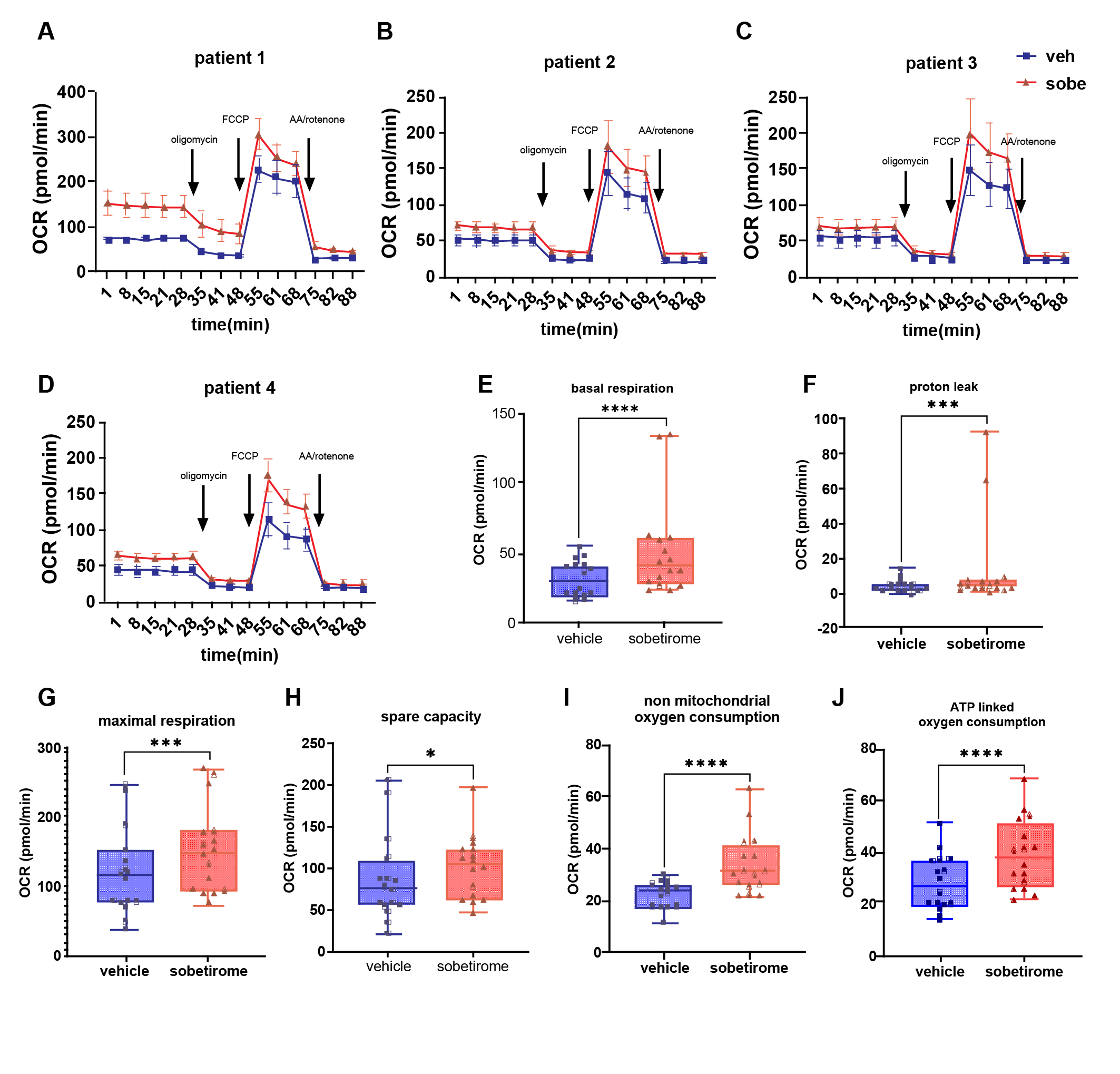
**Supplementary Figure S6.**

**Supplementary Figure S6. IPF-patient-derived fibroblast mitochondrial function is improved by sobetirome**. **(A-D)** The oxygen consumption rate was measured using the Seahorse device in cells derived from 4 different patients. **(E-H)** Indicated parameters were calculated. n=4 for A-D; Student’s t-test was used for F and I, and the Wilcoxon signed rank test was used for E, G, H, and J. *=p<0.05, ***=p<0.001, ****=p<0.0001

**Supplementary Figure S7.**

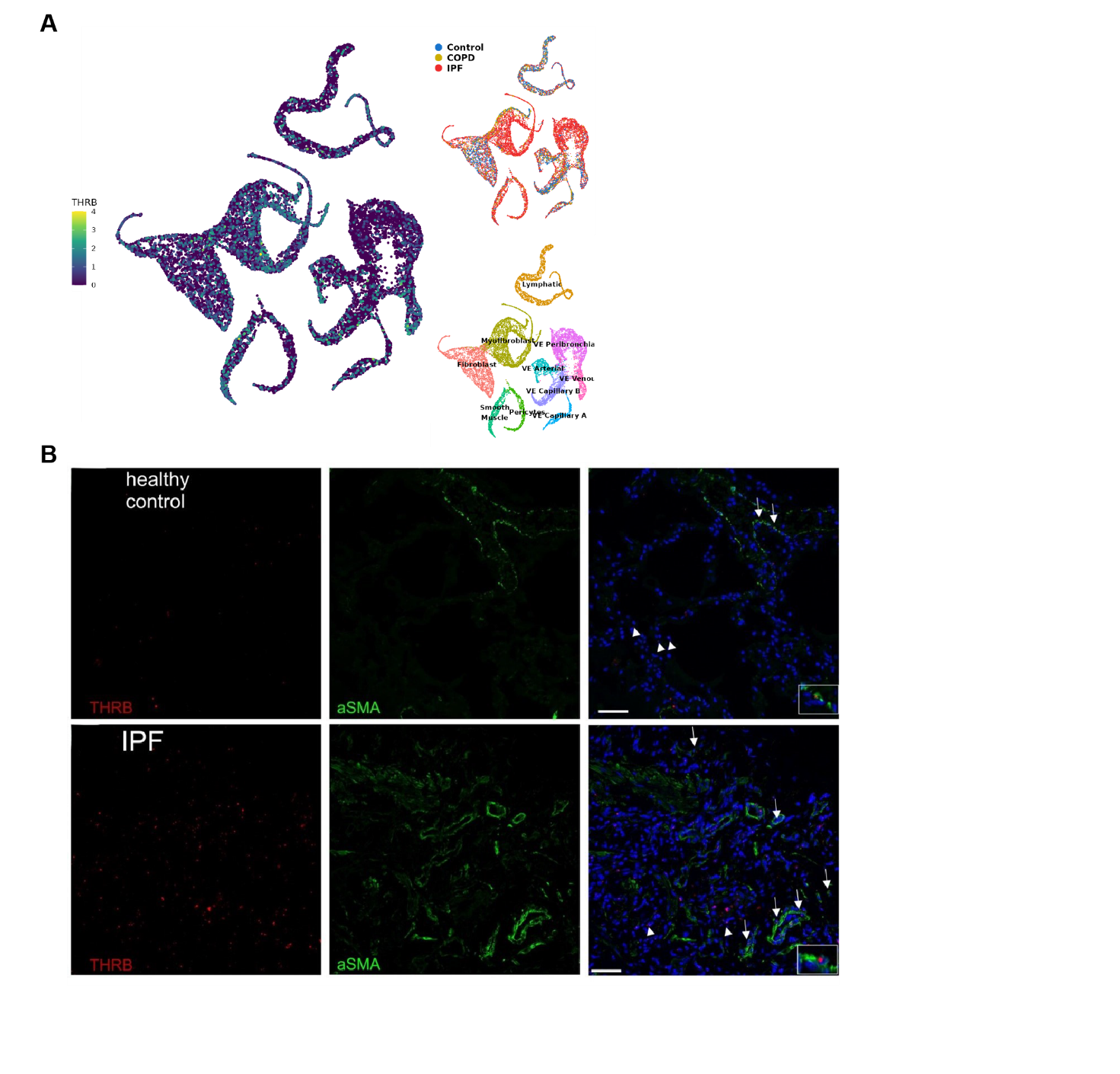

**Supplementary Figure S7. THRB is expressed in aSMA-positive cells in IPF patients.** **(A)** Analysis of a previously published single-cell RNASeq dataset [1] revealed expression of THRB in alveolar fibroblasts/myofibroblasts**. (B, C)** THRB (red) was detected by in situ hybridization, and aSMA (green) was stained by IF. Insets and arrows show double-positive cells; arrowheads depict cells negative for aSMA but positive for THRB.

**Supplementary Figure S8.**
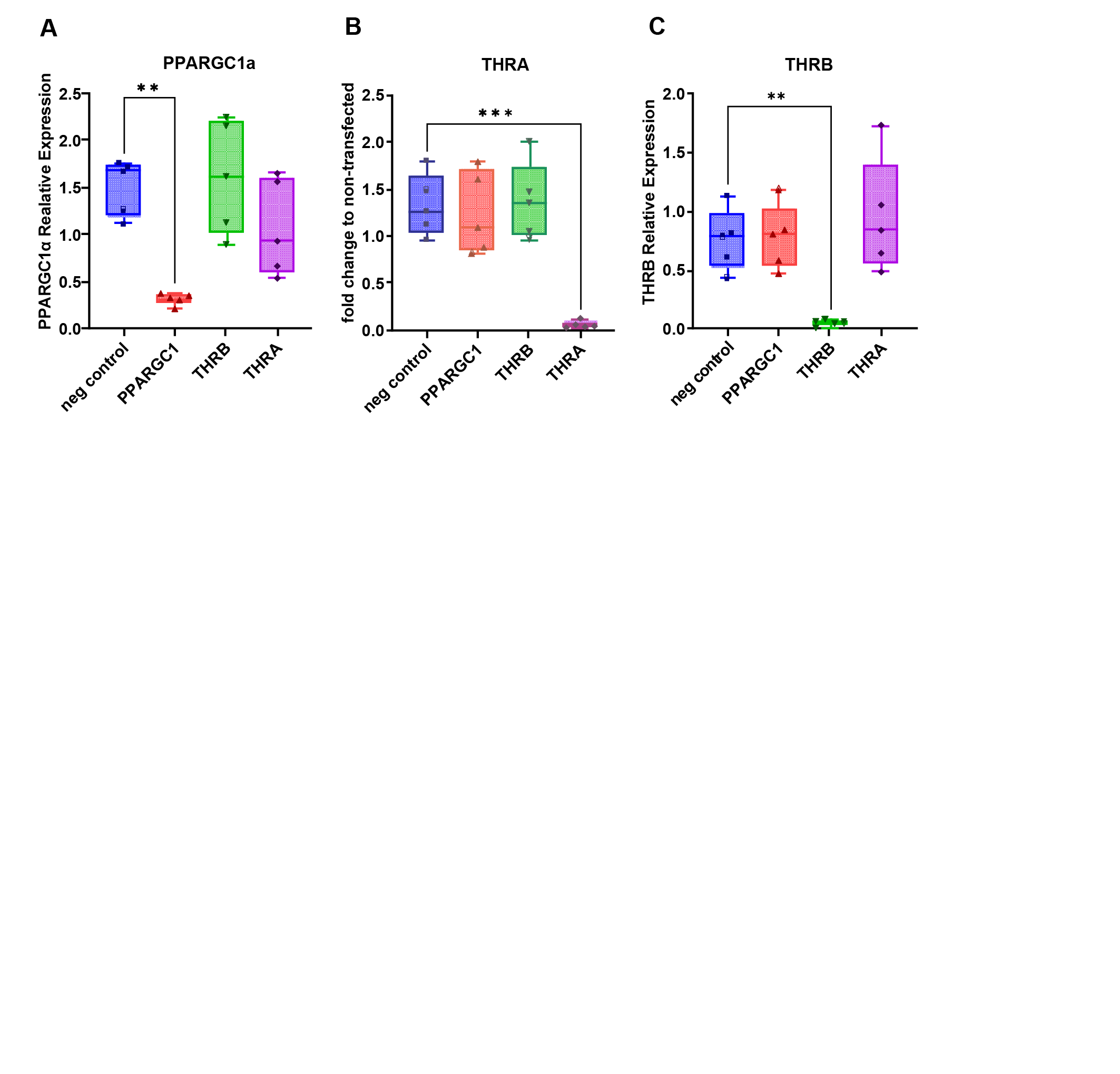

**Supplementary Figure S8. IPF Fibroblasts were transfected twice with either negative control siRNA (neg control) or siRNA targeting *PPARGC1α*, *THRA,* or *THRB****.* The levels of **(A)** PPARGC1α, **(B)** THRA, and **(C)** THRB were assessed after 7 days and are provided as normalized to a non-transfected control. Analysis was performed by One-way ANOVA. **=p<0.01, ***=p<0.001.

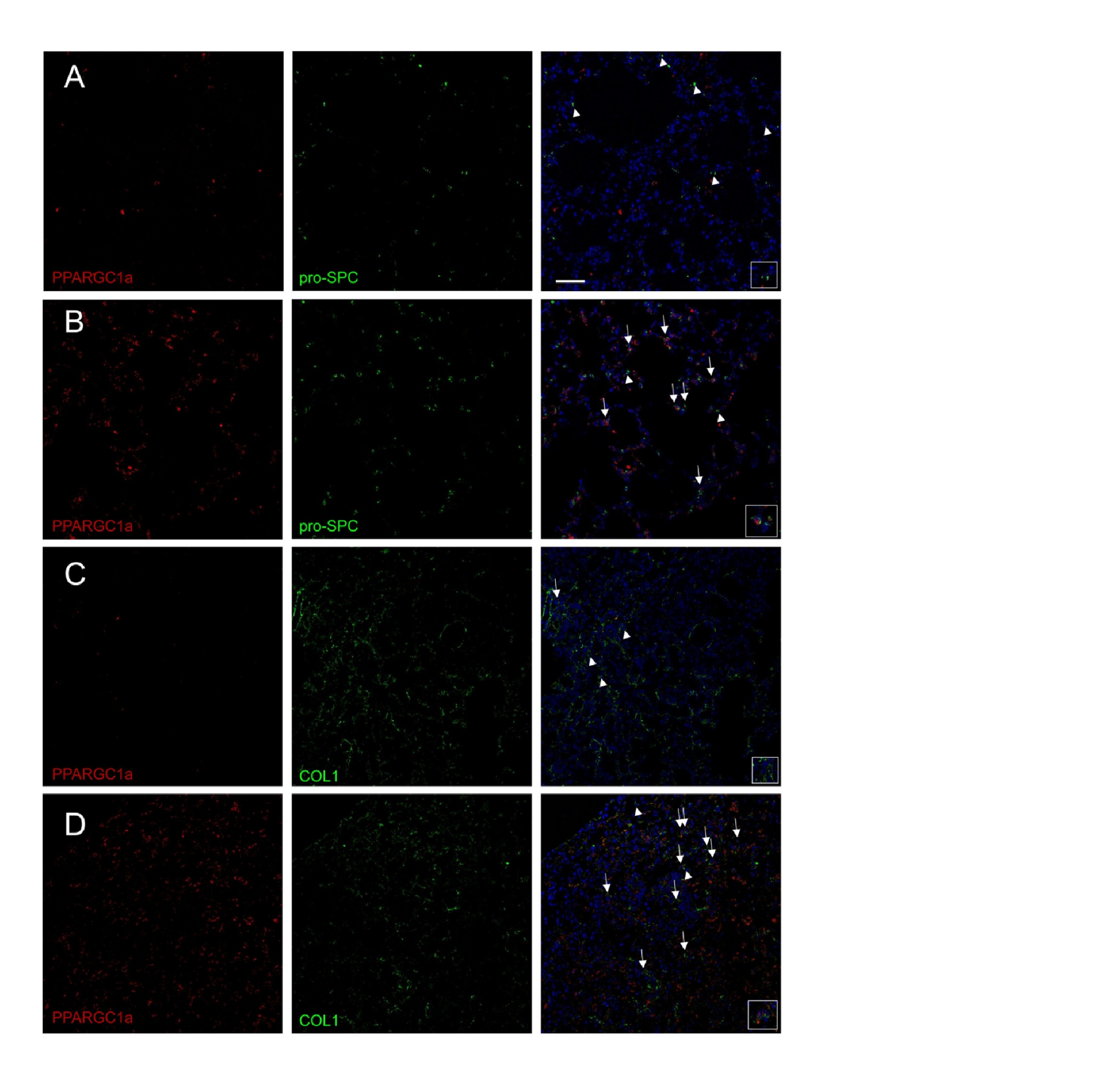
**Supplementary Figure S9.**

**Supplementary Figure S9. Sobetirome treatment increases PPARGC1a staining and nuclear localization in bleomycin-induced fibrosis.** Sections from mice receiving **(A** and C) bleomycin+ vehicle or **(B** and **D)** bleomycin + sobetirome treatment were stained with antibodies directed against (red) PPARGC1a and (A and B, green) pro-SPC or (C and D, green) Col1. Insets show single cells (A and C) without or (B and D) with nuclear localization of PPARGC1a. Arrows show double-positive cells, while arrowheads show cells staining positive for (A and B) pro-SPC or (C and D) Col1. Scale bar is 20µm.

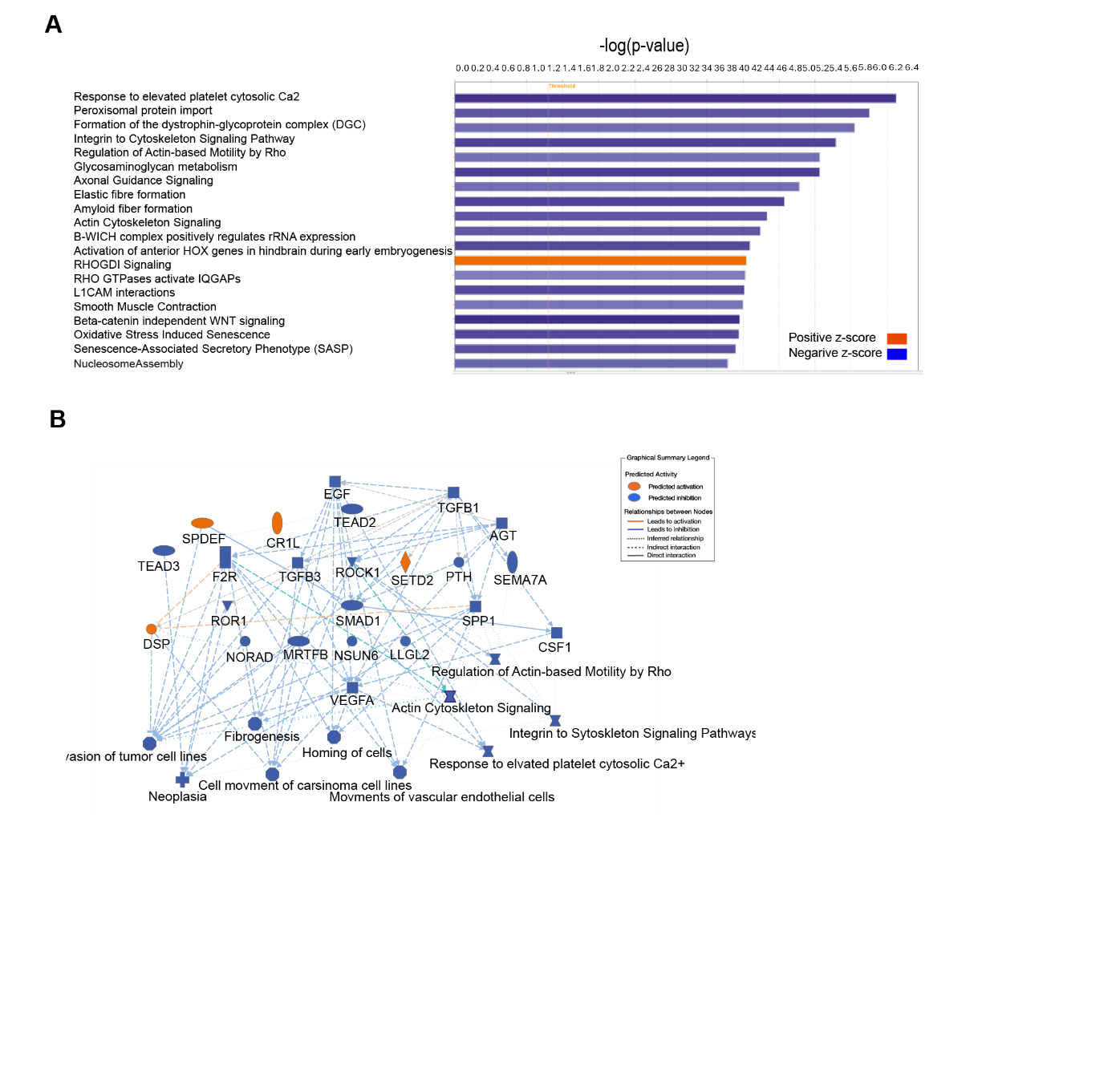
**Supplementary Figure S10.**

**Supplementary Figure S10. Integrative pathway analysis of IPF hPCLS RNA-seq data after sobetirome treatment. (A)** Canonical pathway enrichment analysis was generated using Ingenuity Pathway Analysis (IPA). The pathway chart displays significantly enriched canonical pathways identified from down-regulated differential gene expression in human precision-cut lung slices (hPCLS), ranked by statistical significance (−log10 *p*-value). Bars represent enrichment magnitude, and color coding indicates predicted pathway activation states based on z-scores. **(B)** Summary of the integrative analysis framework combining significantly enriched canonical pathways and predicted upstream regulators. This integrative framework combines significantly enriched canonical pathways, upstream regulators, and disease/functions (p ≤ 0.05, with regulators and functions also meeting |z|-score ≥ 2).

**Supplementary Tables**

**Supplementary Table S1: Top 100 regulated genes affected by sobetirome in mice after bleomycin treatment. (p-value based)**

| **gene** | **baseMean** | **log2FoldChange** | **pvalue** | **padj** |
| --- | --- | --- | --- | --- |
| Sulf1 | 2423.54791 | -2.333032669 | 4.02E-30 | 8.37E-26 |
| Ikbip | 1190.73681 | -0.990040989 | 4.85E-26 | 5.05E-22 |
| Col5a1 | 7188.43721 | -2.45827041 | 8.89E-26 | 6.17E-22 |
| Ccl5 | 341.335639 | 1.872261685 | 1.53E-25 | 7.98E-22 |
| Col1a1 | 32762.2771 | -2.906295407 | 2.59E-24 | 9.28E-21 |
| Fkbp10 | 1035.42584 | -1.740734332 | 2.67E-24 | 9.28E-21 |
| Nlgn2 | 2065.80501 | -1.264797635 | 1.28E-23 | 3.81E-20 |
| Lrrc15 | 64.0384015 | -4.86703916 | 5.55E-23 | 1.44E-19 |
| Col5a2 | 10628.9826 | -2.554321963 | 7.29E-23 | 1.69E-19 |
| Slc9a1 | 1870.85401 | -0.695416267 | 1.59E-22 | 3.32E-19 |
| Cthrc1 | 281.336804 | -2.602369909 | 2.47E-22 | 4.67E-19 |
| Colq | 671.498926 | 3.567214304 | 3.62E-22 | 6.27E-19 |
| Baiap2l1 | 1223.37553 | 1.134354022 | 4.04E-22 | 6.47E-19 |
| Timp1 | 921.938026 | -3.71853467 | 7.09E-22 | 1.05E-18 |
| Fn1 | 69140.0119 | -2.874251991 | 1.20E-21 | 1.66E-18 |
| Serpina3n | 7360.31179 | -2.794216676 | 1.33E-21 | 1.73E-18 |
| Col1a2 | 34958.1543 | -2.395952485 | 2.90E-21 | 3.56E-18 |
| Fstl1 | 9792.60625 | -2.190726898 | 9.37E-21 | 1.08E-17 |
| Fmn1 | 474.963597 | -1.358018797 | 1.63E-20 | 1.79E-17 |
| Calu | 11783.1526 | -0.838826416 | 3.13E-20 | 3.16E-17 |
| Zfp521 | 782.304546 | -1.276700342 | 3.19E-20 | 3.16E-17 |
| Gria1 | 608.257886 | 3.273573993 | 5.36E-20 | 5.07E-17 |
| Tnfrsf11b | 209.670372 | -1.984792942 | 1.07E-19 | 9.66E-17 |
| Crtap | 1246.52234 | -0.921151205 | 1.12E-19 | 9.76E-17 |
| Pex6 | 1113.58736 | 0.743564415 | 1.24E-19 | 1.03E-16 |
| Sfrp1 | 2607.33803 | -3.231144056 | 2.63E-19 | 2.11E-16 |
| Zfp469 | 483.397121 | -1.94819098 | 1.37E-18 | 1.06E-15 |
| Ccl7 | 74.1670103 | -5.759005881 | 3.22E-18 | 2.40E-15 |
| Fbn1 | 12394.6951 | -1.90796883 | 3.34E-18 | 2.40E-15 |
| Tppp3 | 5423.01507 | 1.68298088 | 6.38E-18 | 4.43E-15 |
| Gm33248 | 153.198118 | 3.481240442 | 6.76E-18 | 4.54E-15 |
| Fzd1 | 2791.7851 | -1.070731559 | 7.85E-18 | 5.11E-15 |
| Adam12 | 418.642857 | -3.890723016 | 8.93E-18 | 5.63E-15 |
| Sema3d | 908.832322 | -2.118300261 | 1.78E-17 | 1.09E-14 |
| Srpx2 | 450.646357 | -1.883213642 | 2.61E-17 | 1.55E-14 |
| Dab2 | 3647.26851 | -1.561602231 | 3.79E-17 | 2.19E-14 |
| Ppp1r14c | 2012.25697 | 1.573833289 | 5.38E-17 | 3.03E-14 |
| Igfbp4 | 9036.40776 | -1.554249052 | 5.74E-17 | 3.15E-14 |
| Baiap3 | 333.541714 | 1.645906053 | 6.19E-17 | 3.30E-14 |
| Cavin3 | 762.802274 | -0.935849607 | 6.36E-17 | 3.31E-14 |
| Col16a1 | 2085.65975 | -1.309524233 | 8.51E-17 | 4.32E-14 |
| Lox | 5925.50697 | -2.318900158 | 9.12E-17 | 4.52E-14 |
| Klre1 | 141.761912 | 3.310635454 | 1.20E-16 | 5.82E-14 |
| Gpt | 628.583587 | 1.318380596 | 1.27E-16 | 6.01E-14 |
| Mgp | 28365.5865 | -1.641871959 | 1.36E-16 | 6.31E-14 |
| Emilin1 | 3129.29853 | -1.069298499 | 1.54E-16 | 6.95E-14 |
| Tnfrsf1a | 4983.72715 | -0.597239756 | 1.88E-16 | 8.35E-14 |
| P3h1 | 916.590497 | -1.140335599 | 2.38E-16 | 1.03E-13 |
| Ckap4 | 2412.90966 | -1.242402689 | 2.50E-16 | 1.06E-13 |
| Grk3 | 3595.21816 | 0.755488677 | 2.73E-16 | 1.14E-13 |
| Cd226 | 298.7544 | 1.628282813 | 3.13E-16 | 1.28E-13 |
| Slc35f6 | 1178.65501 | -0.656207017 | 3.37E-16 | 1.35E-13 |
| Mettl7a1 | 13877.6407 | 1.546049214 | 4.72E-16 | 1.85E-13 |
| Ctsl | 7334.82307 | -1.059899387 | 5.26E-16 | 2.03E-13 |
| Dse | 1326.52958 | -0.719138829 | 6.75E-16 | 2.56E-13 |
| Sdk1 | 319.541748 | -1.402407842 | 9.61E-16 | 3.57E-13 |
| Nkg7 | 193.434147 | 2.54104072 | 1.57E-15 | 5.75E-13 |
| Acss1 | 2156.68041 | 0.980098785 | 1.85E-15 | 6.64E-13 |
| Acoxl | 1206.40912 | 3.170701974 | 1.95E-15 | 6.87E-13 |
| Serpine2 | 5779.99279 | -1.767512036 | 2.19E-15 | 7.61E-13 |
| Klri2 | 139.572667 | 2.740174488 | 2.39E-15 | 8.16E-13 |
| Maob | 1345.85522 | 0.970703359 | 2.51E-15 | 8.44E-13 |
| Loxl3 | 618.991351 | -1.631795965 | 2.87E-15 | 9.49E-13 |
| Ak5 | 60.6974028 | -3.098564819 | 3.20E-15 | 1.04E-12 |
| Sec61a1 | 4072.09289 | -0.498784496 | 3.36E-15 | 1.08E-12 |
| Vim | 25574.8622 | -0.62283753 | 4.49E-15 | 1.41E-12 |
| Txlng | 1314.00412 | 0.817435797 | 4.68E-15 | 1.45E-12 |
| Rac3 | 776.176508 | 1.31059042 | 5.79E-15 | 1.77E-12 |
| Adamts12 | 992.235065 | -3.830506087 | 6.17E-15 | 1.86E-12 |
| Pappa | 326.55588 | -2.879886285 | 6.43E-15 | 1.91E-12 |
| Adam15 | 2573.15865 | -0.842164897 | 7.36E-15 | 2.16E-12 |
| Klrc1 | 89.4977568 | 2.452043341 | 9.35E-15 | 2.70E-12 |
| Tnfrsf23 | 172.25421 | -2.090134525 | 9.55E-15 | 2.72E-12 |
| Acot1 | 552.508537 | 1.916960771 | 9.91E-15 | 2.79E-12 |
| Inhba | 486.873354 | -4.026659992 | 1.04E-14 | 2.90E-12 |
| Cecr6 | 98.3467523 | 3.555246627 | 1.06E-14 | 2.91E-12 |
| Aldh1a1 | 25243.1275 | 1.169329142 | 1.08E-14 | 2.91E-12 |
| Adam22 | 648.040126 | 1.026298532 | 1.13E-14 | 3.02E-12 |
| Arhgef38 | 1633.72911 | 1.971834304 | 1.15E-14 | 3.03E-12 |
| Cat | 8531.78017 | 1.086803161 | 1.16E-14 | 3.03E-12 |
| Myo7a | 1534.15822 | -1.215980549 | 1.26E-14 | 3.23E-12 |
| Inf2 | 2095.66398 | -0.982587472 | 1.37E-14 | 3.49E-12 |
| Adamts7 | 207.429415 | -1.860210351 | 1.41E-14 | 3.54E-12 |
| Loxl2 | 3067.02986 | -2.288022221 | 1.43E-14 | 3.55E-12 |
| Col8a1 | 1157.64682 | -2.359921757 | 1.55E-14 | 3.80E-12 |
| Lingo1 | 74.7973416 | -2.653344205 | 1.62E-14 | 3.93E-12 |
| Kirrel | 1976.37623 | -1.155172063 | 1.75E-14 | 4.19E-12 |
| Dnajc10 | 3852.33393 | -0.49226655 | 2.28E-14 | 5.40E-12 |
| Banp | 633.65026 | 0.893516993 | 2.59E-14 | 6.06E-12 |
| Sftpc | 207676.611 | 1.670819726 | 3.16E-14 | 7.31E-12 |
| Grb14 | 1468.64112 | 1.013384539 | 3.83E-14 | 8.76E-12 |
| Enpp1 | 544.43011 | -1.583534264 | 4.14E-14 | 9.38E-12 |
| Fmod | 1445.35327 | -3.333193606 | 4.23E-14 | 9.48E-12 |
| Fabp12 | 60.3524681 | 3.624852147 | 4.51E-14 | 9.98E-12 |
| Cbr2 | 50293.8527 | 1.69219296 | 5.11E-14 | 1.12E-11 |
| Cspg4 | 1438.78988 | -1.117105155 | 5.87E-14 | 1.27E-11 |
| Tfdp1 | 1194.43724 | -0.794714508 | 6.21E-14 | 1.33E-11 |
| Mmp14 | 4708.87411 | -1.947432144 | 6.51E-14 | 1.38E-11 |
| Cst8 | 74.1304129 | 3.88012916 | 6.58E-14 | 1.38E-11 |

**Supplementary Table S2: Top 100 regulated genes affected by sobetirome in IPF lung. (p-value based)**

| **gene** | **log2FoldChange** | **pvalue** | **padj** |
| --- | --- | --- | --- |
| GJA5 | -3.426356837 | 7.12E-07 | 0.001821862 |
| LINC00578 | -2.853782762 | 0.01535933 | 0.254215912 |
| AC092691.1 | -2.817989947 | 0.0062129 | 0.189309652 |
| NAV2-AS4 | -2.783127355 | 0.00042465 | 0.056166878 |
| WISP2 | -2.751334123 | 0.00020713 | 0.038648299 |
| MEOX2 | -2.623933348 | 0.00047799 | 0.059732026 |
| MT-RNR1 | -2.550238687 | 0.02415959 | 0.301366361 |
| MFAP4 | -2.510412461 | 5.01E-05 | 0.021982683 |
| HMGCLL1 | -2.49532526 | 7.94E-05 | 0.025369393 |
| AC026355.1 | -2.479595746 | 0.03200745 | 0.333880042 |
| NAP1L3 | -2.477115128 | 0.01007647 | 0.224903674 |
| IGSF10 | -2.457441003 | 0.01659593 | 0.263855813 |
| MEST | -2.441066435 | 7.75E-05 | 0.025308953 |
| PODN | -2.410662084 | 0.00011593 | 0.02964566 |
| ADH1B | -2.385112357 | 0.0145005 | 0.24946969 |
| ALPK2 | -2.377289045 | 0.0052982 | 0.179943549 |
| RN7SL471P | -2.376289694 | 0.00011259 | 0.02964566 |
| BMP5 | -2.352416931 | 0.02978392 | 0.323540922 |
| CCDC80 | -2.263028121 | 0.01464982 | 0.250303158 |
| OLFML1 | -2.239375559 | 0.00014857 | 0.032147208 |
| FIBIN | -2.197410068 | 0.00040607 | 0.054224537 |
| NCAM2 | -2.191761772 | 0.0413741 | 0.36315953 |
| MT-RNR2 | -2.186303744 | 0.0352774 | 0.347248121 |
| ISLR | -2.184271982 | 0.00138746 | 0.10590953 |
| GPR1 | -2.160604411 | 0.01012875 | 0.224903674 |
| CGREF1 | -2.149199546 | 0.00031707 | 0.046777642 |
| DLG2 | -2.147062679 | 0.00044598 | 0.057989221 |
| POSTN | -2.107057853 | 1.82E-05 | 0.015893683 |
| TMEM35A | -2.085586368 | 0.00120528 | 0.097844652 |
| HSPB6 | -2.068346488 | 0.01440106 | 0.24946969 |
| BEND6 | -2.058959119 | 0.00022525 | 0.039604002 |
| PLA2G5 | -2.055744465 | 0.01372525 | 0.246442838 |
| F10 | -2.012513074 | 0.002428 | 0.135098669 |
| GLT8D2 | -1.987129146 | 0.00019783 | 0.038398211 |
| RARRES1 | -1.979845299 | 0.00016562 | 0.035089282 |
| ALPL | -1.97852755 | 0.00027641 | 0.043721689 |
| SFRP4 | -1.973725727 | 0.00034335 | 0.049697607 |
| CXCL12 | -1.942071017 | 0.01598402 | 0.258629308 |
| CTSK | -1.933970706 | 2.49E-05 | 0.018142051 |
| LRRC17 | -1.933210361 | 0.01875022 | 0.276309058 |
| SAMD3 | -1.927724503 | 0.02300988 | 0.294878061 |
| ADAMTS16 | -1.923648016 | 0.00020021 | 0.038398211 |
| SMPDL3A | -1.909404305 | 0.00147188 | 0.109096561 |
| PTGDS | -1.907815004 | 4.24E-05 | 0.021748356 |
| LRRN4CL | -1.887251376 | 0.00681743 | 0.196986429 |
| OSR1 | -1.881337868 | 0.00150755 | 0.110671205 |
| ANK2 | -1.879153982 | 0.02252036 | 0.293256952 |
| TMEM119 | -1.866120551 | 0.00073195 | 0.074978634 |
| FGF14-IT1 | -1.863031067 | 0.01302207 | 0.242767379 |
| HOXB-AS3 | -1.843177603 | 0.00166396 | 0.114463884 |
| SPON2 | -1.838819454 | 5.25E-05 | 0.022380249 |
| PQLC2L | -1.834791509 | 0.00058952 | 0.066324367 |
| VEPH1 | -1.830657421 | 0.0056133 | 0.184027321 |
| OMG | -1.829850448 | 0.00043829 | 0.05747592 |
| ITIH5 | -1.818866186 | 0.00018664 | 0.037679651 |
| KCTD16 | -1.815548515 | 0.01056173 | 0.227965602 |
| SELENOP | -1.797471612 | 0.02427704 | 0.302237924 |
| CD248 | -1.786170432 | 0.00574399 | 0.185379756 |
| CASS4 | -1.782692139 | 0.00702079 | 0.199481553 |
| GDF10 | -1.764573651 | 0.00977585 | 0.223533309 |
| C1QTNF2 | -1.752689888 | 0.01732387 | 0.268484943 |
| ACOX2 | -1.749573873 | 0.00011558 | 0.02964566 |
| SLC1A7 | -1.747220445 | 0.00835094 | 0.213197221 |
| RCAN2 | -1.736323284 | 0.00715001 | 0.200570853 |
| PRELP | -1.724971262 | 0.00247183 | 0.135098669 |
| FAM19A5 | -1.717653189 | 0.00286514 | 0.142259908 |
| DOK5 | -1.714227915 | 0.00168964 | 0.114486818 |
| DCN | -1.696802166 | 0.00067845 | 0.072288016 |
| LINC01936 | -1.681322226 | 0.02577426 | 0.309750352 |
| C16orf89 | -1.669488415 | 0.01970892 | 0.27904962 |
| KCND3 | -1.664238767 | 0.00010194 | 0.028799427 |
| SAR1AP2 | -1.663220851 | 0.00055593 | 0.064133062 |
| AC099066.2 | -1.660165938 | 0.010717 | 0.227965602 |
| PCOLCE | -1.658269013 | 0.00089909 | 0.081625875 |
| PTX3 | -1.648598995 | 0.00398912 | 0.160643212 |
| HNRNPA1P29 | -1.642769132 | 0.00185706 | 0.12022341 |
| OLFML3 | -1.634849058 | 0.00105837 | 0.090718576 |
| PLAC9 | -1.629421787 | 0.00075608 | 0.074978634 |
| PTN | -1.624317416 | 0.0260639 | 0.311635979 |
| NAV2-AS5 | -1.617463672 | 0.01912589 | 0.278367684 |
| CD36 | -1.600073452 | 0.00545087 | 0.18208081 |
| SGCD | -1.59599309 | 0.01005209 | 0.224903674 |
| PDGFRL | -1.577282937 | 0.00876452 | 0.214249933 |
| CHI3L2 | -1.573783166 | 0.00020836 | 0.038648299 |
| SCN7A | -1.570708112 | 0.01476252 | 0.250951925 |
| RPS12P5 | -1.56418184 | 0.00278871 | 0.141916998 |
| ACTA2 | -1.560018375 | 0.0037411 | 0.153886623 |
| GLIPR1 | -1.552475757 | 0.00626262 | 0.189310308 |
| PIEZO2 | -1.547047732 | 0.00464366 | 0.170449149 |
| PKDCC | -1.546344942 | 0.00555023 | 0.183133861 |
| HIST1H1A | -1.536581355 | 0.0005041 | 0.060901379 |
| ECM2 | -1.532328605 | 0.00362634 | 0.151690511 |
| SFRP2 | -1.517693267 | 6.51E-06 | 0.006681448 |
| PCDH9 | -1.517487596 | 0.0286423 | 0.321405742 |
| RGS4 | -1.507636823 | 0.00364942 | 0.15215519 |
| HIST1H2AJ | -1.507494959 | 0.00324589 | 0.146475532 |
| DIO3OS | -1.507258398 | 0.01674282 | 0.264830018 |
| AC011389.1 | -1.501471482 | 0.02152208 | 0.290128167 |
| SMIM10 | -1.488777882 | 0.01397461 | 0.248738408 |

**Supplementary Table S3. List of antibodies.**

|  | **antibody** | **host species** | **company** | **number** | **IF/ICH/FACS** | **WB** |
| --- | --- | --- | --- | --- | --- | --- |
| **primary AB** | pro-SP-C | rabbit | millipore | AB3786 | 1:500 | - |
|  | beta-actin | mouse | Santa Cruz | sc-47778 | - | 1:1000 |
|  | PINK1 | rabbit | abcam | ab23707 | - | 1:1000 |
|  | aSMA | rabbit | Abcam | 5694 | 1:500 | - |
|  | Collagen I | rabbit | abcam | ab34710 | 1:100 | - |
|  | Caspase 3 | rabbit | LSBio | LS‑C351924 | - | 1:1000 |
|  | Cleaved Caspase-3 | rabbit | Cell signaling | 9661S | - | 1:1000 |
|  | pAMPK-alpha | rabbit | Cell signaling | 2535S | - | 1:1000 |
|  | AMPK-alpha | rabbit | Cell signaling | 2630S | - | 1:1000 |
|  | PPARGC1alpha | rabbit | Novus | NBP1-04676 | 1:100 | 1:1000 |
|  | PPARGC1alpha | rabbit | abcam | ab54481 | - | 1:1000 |
|  | CD16/32 | rat | biolegend | 101306 | 1:100 | - |
|  | CD45 | rat | biolegend | 103108 | 1:100 | - |
|  | CD11b | rat | biolegend | 101206 | 1:100 | - |
|  | CD11c | rat | biolegend | 117306 | 1:100 | - |
|  | F4/80 | rat | biolegend | 123108 | 1:100 | - |
|  | CD19 | rat | biolegend | 152404 | 1:100 | - |
| **Secondary AB** | anti rabbit AF594 | goat | Invitrogen | A-11037 | 1:500 | - |
|  | anti rabbit AF488 | goat | Invitrogen | A-11008 | 1:500 | - |
|  | anti rabbit AF647 | goat | JacksonIR | 111-605-144 | 1:500 | - |
|  | anti rabbit HRP | donkey | Amersham | NA934V | - | 1:2500 |

| **target gene** | **company** | **number** | **species** |
| --- | --- | --- | --- |
| PPARGC1 | Thermo Fisher Scientific | Mm01208835_m1 | Mouse |
| gusb | Thermo Fisher Scientific | Mm01197698_m1 | Mouse |
| gapdh | Thermo Fisher Scientific | Mm99999915_g2 | Mouse |
| col1a1 | Thermo Fisher Scientific | Mm01302043_g1 | Mouse |
| PPARGC1 | Thermo Fisher Scientific | Hs00173304_m1 | Human |
| COL1A1 | Thermo Fisher Scientific | Hs00173304_m1 | Human |
| THRA | Thermo Fisher Scientific | Hs00230861_m1 | Human |
| THRB | Thermo Fisher Scientific | Hs00268470_m2 | Human |

**Supplementary Table S4: List of primers.**

1.	Adams, T.S. et al., *Single-cell RNA-seq reveals ectopic and aberrant lung-resident cell populations in idiopathic pulmonary fibrosis.* Sci Adv, 2020. **6**(28): p. eaba1983.
